# Unexpected diversity of *Isidoides* (Anthozoa: Octocorallia: Isidoidae) revealed by morphology and phylogenomic analysis with descriptions of three new species

**DOI:** 10.1101/2025.09.11.675584

**Authors:** Yu Xu, Jaret P. Bilewitch, Eric Pante, Zifeng Zhan, Sadie Mills, Malcolm R. Clark, Kuidong Xu

**Affiliations:** Laboratory of Marine Organism Taxonomy and Phylogeny, Shandong Province Key Laboratory of Marine Biodiversity and Bio-resource Sustainable Utilization, Institute of Oceanology, Chinese Academy of Sciences, Qingdao 266071, China; New Zealand Institute of Earth Science Ltd (NZIES), 301 Evans Bay Parade, Wellington 6021, New Zealand; Univ Brest, CNRS, IRD, Ifremer, UMR 6539, LEMAR, Plouzané, France; Institut Systématique Evolution Biodiversité (ISYEB), Muséum national d’Histoire naturelle, CNRS, Sorbonne Université, EPHE, Université des Antilles, 43 rue Cuvier, CP 26, 75005 Paris, France; University of Chinese Academy of Sciences, Beijing 100049, China

**Keywords:** deep sea, phylogeny, Scleralcyonacea, seamount, taxonomy, ultraconserved elements, species delimitation

## Abstract

Isidoidae Heestand Saucier, France & Watling, 2021 is a rare octocoral family currently represented by a single genus and species, *Isidoides armata* Nutting, 1910 recorded in the western Pacific Ocean. The taxonomic status and diversity of *Isidoides* is unclear, due to the lack of diagnostic taxonomic features and limited taxon sampling. Based on 23 *Isidoides* specimens obtained from the northwestern to southwestern Pacific, we carried out morphological and phylogenetic analyses to reveal the taxonomic status of new species and develop reliable features for species identification. The 23 specimens could be classified into four well-supported clades by the phylogenomic analysis of ultraconserved elements (UCEs), four groups by *28S rDNA*, and two groups by *mtMutS-cox1*. Integrating morphology and molecular data, we uncovered unexpected diversity of *Isidoides* composed of the known species *Isidoides armata* and three new species, viz., *I. elegans* sp. nov., *I. gracilis* sp. nov. and *I. pseudarmata* sp. nov. The morphological analysis showed high intraspecific morphological variation in colony color and the size, shape and arrangement of polyps. By contrast, sclerite forms with their surface sculpturing are more diagnostic features for species identification. Our phylogenetic and species delimitation analyses indicate that UCEs have higher resolution than the nuclear *28S rDNA* and the mitochondrial genes *mtMutS* and *cox1* for species discrimination within Isidoidae.

ZooBank: urn:lsid:zoobank.org:pub:392485F5-502E-4383-B153-45B167571190

## Introduction

Octocorals are often the most conspicuous megafauna in deep-sea benthic ecosystems with hard substrate, such as seamounts where they may form coral forests and serve as vital habitats for marine life (e.g., Krieger and Wing, 2002; Buhl-Mortensen and Mortensen, 2005; Parimbelli, 2020; Rossi et al., 2022; Morrissey et al., 2023). With their significant ecological value and physical fragility, octocorals have been considered an indicator group for the conservation of vulnerable marine ecosystems, particularly seamount ecosystems (Dautova, 2019; Watling and Auster, 2021). Historically, octocoral taxonomy and systematics have relied exclusively on morphological traits (e.g., Bayer, 1981). Modern advances in molecular methods have prompted major revisions to their classification (McFadden et al., 2022). Nevertheless, challenges in octocoral systematics persist, and the diversity and relationships of many groups are still largely unknown (Kessel et al., 2022).

Isidoidae Heestand Saucier, France & Watling, 2021 is a family of rare octocorals comprising only the monotypic genus *Isidoides* Nutting, 1910, with the species *I. armata* Nutting, 1910 recorded from the tropical western Pacific Ocean (Nutting, 1910). The genus has been assigned to various families, including Chrysogorgiidae, Ellisellidae, Primnoidae and Isididae, due to its conflicting morphological features (Bayer, 1979; Bayer and Stefani, 1988; Bayer and Grasshoff, 1994). The molecular phylogenetic study of Pante et al. (2013) showed a close affinity between *Isidoides* and the Isididae subfamily Keratoisidinae (now elevated to family Keratoisididae) but placed the genus as *incertae sedis*. Later, the family Isidoidae was established to contain the genus *Isidoides* with its non-articulated axis and a distinct sclerite form, based on the phylogenetic analysis of *mtMutS* + *18S rDNA* and a unique arrangement of mitochondrial genes (Heestand Saucier et al., 2021). The phylogenomic analysis of ultraconserved elements (UCEs) corroborated the validity and distinctiveness of the family Isidoidae, supporting a sister relationship with Keratoisididae (e.g., McFadden et al., 2021, 2022).

When re-examining the original collections, Pante et al. (2013) found that the holotype (a fragment) of *I. armata* illustrated in Nutting (1910) was lost but another fragment from a different locality was available. To avoid future taxonomic confusion, Pante et al. (2013) designated Nutting’s illustrated (but lost) fragment as the lectotype, and the existing fragment as a paralectotype. Nonetheless, they could not confirm whether the holotype and the alternative fragment originated from the same colony. Meanwhile, they succeeded in DNA-sequencing three *mtMutS* haplotypes (A, B and C) from newly collected specimens from New Caledonia and New Zealand. Pante et al. (2013) suggested that the paralectotype of *I. armata* and the specimens representing the three haplotypes could together represent four species. They noted that the paralectotype differs from the New Caledonia and New Zealand specimens in having more slender polyps and a thinner, transparent coenenchyme, and that specimens with haplotype C possess a distinctive dark golden-brown to black axis. However, the available data were insufficient to separate these specimens, which were conservatively treated as *Isidoides armata* (Pante et al., 2013).

The 5’-end of the mitochondrial gene *mtMutS* has been commonly used in the molecular identification of octocorals, and multilocus barcodes (*mtMutS* + *cox1*) can discriminate roughly 70% of octocoral morphospecies (e.g., McFadden et al., 2010, 2011; McFadden and van Ofwegen, 2013). A single mutation at *mtMutS* has been used to separate species (Pante and Watling, 2012; Pante et al., 2015). However, in many cases, both the *mtMutS* and *cox1* loci are too conserved for species delimitation. For example, concatenated *mtMutS-cox1* could not discriminate among the congeners *Chrysogorgia pendula*, *C. acanthella* and *C. cylindrata* (Xu et al., 2023). The nuclear *28S rDNA* gene has shown a higher level of genetic variation among chrysogorgiid species and has been proposed as a potential barcode for octocoral species delimitation (McFadden et al., 2014; Quattrini et al., 2019; Xu et al., 2021a, b; Baena et al., 2024). Recently, ultraconserved elements (UCEs) and exons obtained from target-capture enrichment have offered advantages over traditional Sanger sequencing and other next-generation sequencing methods such as RAD-seq, as UCEs can address a wide range of evolutionary and taxonomic questions across deep to shallow timescales within the Anthozoa (Quattrini et al., 2017; Erickson et al., 2021; Untiedt et al., 2021). More recently, genome skimming, or low-coverage whole-genome sequencing (which produces highly fragmented and gapped assemblies due to low read depth), has enabled cost-effective acquisition of UCE loci, mitogenomes and ribosomal genes and has been used for coral phylogenomic studies to review and support taxonomic revisions and evolutionary history reconstruction (Quattrini et al., 2024). However, UCEs have primarily been used to confirm the relationship between higher-level taxa (e.g., McFadden et al., 2022), and so far have only been used for delimitation and identification of species in a few cases (e.g., Erickson et al., 2021; Untiedt et al., 2021; Bridge et al., 2024; McFadden et al., 2024; Borghi et al., 2026; Samimi-Namin et al., 2026).

In this study, we adopted an integrative lineage-based species concept in which species are recognized as independently evolving metapopulation lineages. We examined the morphology of *Isidoides* specimens collected from the northwestern to the southwestern Pacific and used DNA sequencing analyses of barcode and UCE loci to enrich the taxonomy and diversity of *Isidoides* Nutting, 1910. Based on examination of historical and newly collected specimens, we review and revise the genus *Isidoides*.

## Material and methods

### Specimen collection and morphological examination

Six octocoral specimens were collected in the tropical western Pacific during cruises of the R/V *KeXue* (Science) using the ROV *FaXian* (Discovery) and an electro-hydraulic grab. Of these, four specimens of *I. gracilis* sp. nov. were collected in the South China Sea in 2016, 2018 and 2022, and two specimens of *I. elegans* sp. nov. were collected in 2019 from an unnamed seamount on the Caroline Ridge (Fig. 1). *In-situ* photographs of the specimens were taken by the ROV before sampling, whenever possible. After collection, all specimens were photographed on-board and in the laboratory, prior to preservation in 75% ethanol for further morphological and molecular analyses. Five specimens of *I. pseudarmata* sp. nov. and ten specimens of *I. armata* were sourced from the Muséum national d’Histoire naturelle (Paris, France); Two specimens of *I. armata* were sourced from the ESNZ National Invertebrate Collection (Wellington, New Zealand; formerly NIWA); all of which were collected from New Caledonia to New Zealand (Fig. 1, Table 1).

**Figure 1.**
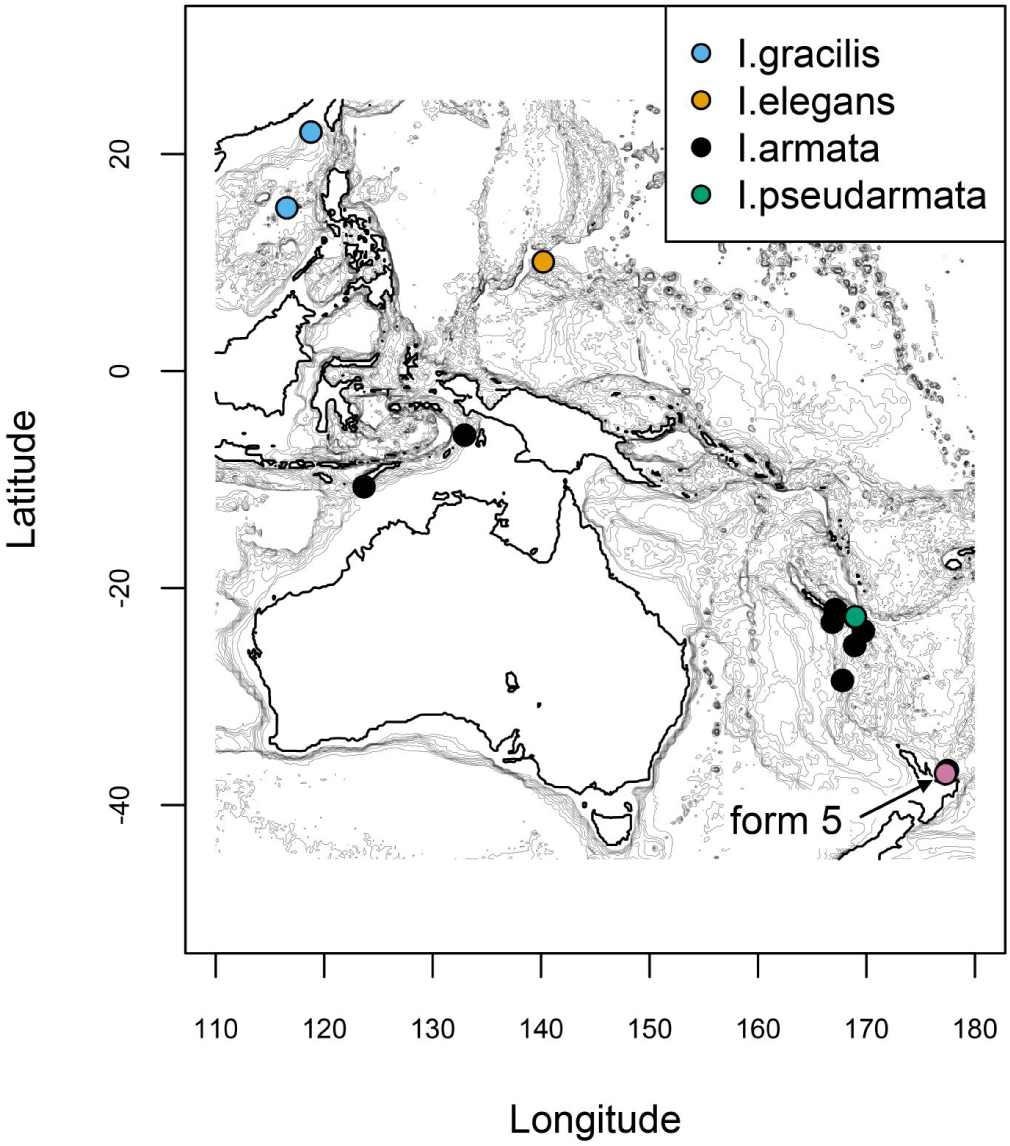
Map showing the collection localities of all discussed *Isidoides* specimens across the Indo-West Pacific and southwest Pacific. Four specimens of *I. gracilis* sp. nov. were collected from the South China Sea; two specimens of *I. elegans* sp. nov. were from an unnamed seamount on the Caroline Ridge; six specimens of *I. pseudarmata* sp. nov. were from New Caledonia; 15 specimens of *I. armata* were recorded, among them, the lectotye from Arafura Sea, the paralectotype from Timor Sea, 11 specimens from New Caledonia to Australia and two specimens from New Zealand. The pink dots represent the collection locality of *I. armata* form 5.

**Table 1.** The sampling localities of *Isidoides* specimens discussed in this study. Some records were obtained from Nutting, 1910 and Pante et al., 2013. “–” = unknown data.

| Species | Collection number or isolated fragment number | Voucher number | *mtMutS* haplotype | Sampling location | Depth (m) | Latitude | Longitude | Collection date |
| --- | --- | --- | --- | --- | --- | --- | --- | --- |
| *Isidoides armata* Nutting, 1910 (lectotype, form 1) | – | – | - | Eastern slope of Elat, Kei Islands, Arafura Sea | 984 | –5.900 | 132.945 | 20/12/1899 |
| *I. elegans* sp. nov. (holotype) | M5043 | MBM286500 | - | An unnamed seamount on the Caroline Ridge | 814 | 10.076 | 140.190 | 31/5/2019 |
| *I. elegans* sp. nov. (paratype) | M5223 | MBM286501 | - | An unnamed seamount on the Caroline Ridge | 962 | 10.078 | 140.203 | 27/5/2019 |
| *I. gracilis* sp. nov. (paratype, form 1) | 11 | MBM286878 | - | Ganquan Plateau, South China Sea | 586–910 | – | – | 14/6/2018 |
| *I. gracilis* sp. nov. (holotype, form 1) | Z076 | MBM286877 | - | Zhenbei Seamount, South China Sea | 797 | 15.049 | 116.558 | 20/7/2022 |
| *I. gracilis* sp. nov. (paratype, form 1) | RY16049 | MBM286879 | - | Northern South China Sea | 811 | 22.035 | 118.776 | –/9/2016 |
| *I. gracilis* sp. nov. (paratype, form 2) | RY16141 | MBM286880 | - | Northern South China Sea | – | 22.035 | 118.776 | –/9/2016 |
| *I. armata* Nutting, 1910 (form 5) | MBM286882 | NIWA163516 | - | Bay of Plenty, New Zealand | 614 | –37.1 | 177.3 | 13/08/1998 |
| *I. armata* Nutting, 1910 (form 4) | MBM286881 | NIWA82680 | - | Bay of Plenty, New Zealand | 878 | –36.81 | 177.47 | 22/04/2012 |
| *I. armata* Nutting, 1910 (form 3) | NORF36/10 | AM G:16995 | A | North of Norfolk Island, Norfolk Ridge, Australia | 1035–1065 | –28.520 | 167.790 | 17/05/2003 |
| *I. armata* Nutting, 1910 (form 3) | NOR220652 | MNHN-IK-2008-1411 | A/C | Athos Seamount, Norfolk Ridge, New Caledonia | 750–800 | –25.267 | 168.933 | 26/10/2003 |
| *I. armata* Nutting, 1910 (form 3) | TER1051 | MNHN-IK-2008-1505 | A | Walpole Island, Loyalty Ridge, New Caledonia | 790–800 | –22.687 | 168.942 | 16/10/2008 |
| *I. armata* Nutting, 1910 (form 3) | TER1012 | MNHN-IK-2008-1504 | A | Walpole Island, Loyalty Ridge, New Caledonia | 720 | –22.675 | 168.962 | 16/10/2008 |
| *I. armata* Nutting, 1910 (form 3) | CP3832-1 | MNHN-IK-2009-1632 | A | South-eastern slope of New Caledonia | 645 | –21.993 | 167.12 | 08/09/2011 |
| *I. armata* Nutting, 1910 (form 3) | TER10084 | MNHN-IK-2008-1507 | B | South-eastern slope of New Caledonia | 500–941 | –21.996 | 167.105 | 16/10/2008 |
| *I. armata* Nutting, 1910 (form 3) | – | MNHN-IK-2008-1506 | - | Loyalty Ridge, New Caledonia | 790–800 | –22.687 | 168.942 | 16/10/2008 |
| *I. armata* Nutting, 1910 (form 3) | TER20516 | MNHN-IK-2008-1510 | B | Mont J, Loyalty Ridge, New Caledonia | 840–800 | –23.958 | 169.722 | 17/10/2008 |
| *I. armata* Nutting, 1910 (form 3) | – | MNHN-IK-2016-2540 | - | South-eastern slope of New Caledonia | 500–941 | –22.000 | 167.100 | 26/10/2008 |
| *I. armata* Nutting, 1910 (form 3) | – | MNHN-IK-2016-2539 | - | Athos Seamount, Norfolk Ridge, New Caledonia | 834–870 | –25.283 | 168.917 | 26/10/2008 |
| *I. armata* Nutting, 1910 (form 3) | – | MNHN-IK-2011-1000 | - | South-eastern slope of New Caledonia | 850 | –23.144 | 166.859 | 29/08/1985 |
| *I. armata* Nutting, 1910 (paralectotype, form 2) | – | ZMA Coel. 02711 | - | South of the western tip of Timor | 520 | –10.65 | 123.67 | – |
| *I. pseudarmata* sp. nov. (paratype) | TER1021 | MNHN-IK-2008-1875 | C | Walpole Island, Loyalty Ridge, New Caledonia | 760–820 | –22.648 | 168.980 | 16/10/2008 |
| *I. pseudarmata* sp. nov. (paratype) | TER1061 | MNHN-IK-2008-1867 | C | Walpole Island, Loyalty Ridge, New Caledonia | 800 | –22.678 | 168.965 | 16/10/2008 |
| *I. pseudarmata* sp. nov. (paratype) | TER1062 | – | A/C | Walpole Island, Loyalty Ridge, New Caledonia | 800 | –22.678 | 168.965 | 16/10/2008 |
| *I. pseudarmata* sp. nov. (paratype) | – | MNHN-IK-2008-1508 | - | Walpole Island, Loyalty Ridge, New Caledonia | 720 | –22.646 | 168.975 | 16/10/2008 |
| *I. pseudarmata* sp. nov. (paratype) | – | MNHN-IK-2008-1509 | - | Walpole Island, Loyalty Ridge, New Caledonia | 800 | –22.678 | 168.965 | 16/10/2008 |
| *I. pseudarmata* sp. nov. (holotype) | – | MNHN-IK-2008-1869 | - | Walpole Island, Loyalty Ridge, New Caledonia | 800 | –22.678 | 168.965 | 16/10/2008 |

Morphological terminology follows Bayer et al. (1983). The general morphology and anatomy were examined using a stereo dissecting microscope. Polyp and branch sclerites were isolated by digesting the tissues in sodium hypochlorite, followed by repeated washing with deionized water. Individual polyps were digested in sodium hypochlorite for 10–30 seconds and gently rinsed with deionized water. The dried polyps and sclerites were mounted on carbon double-adhesive tape, coated, and observed via scanning electron microscopy (SEM) to investigate their structure (Xu et al., 2023). SEM imaging was performed using a Hitachi TM3030Plus SEM, with optimal magnification selected for each sclerite type. Full resolution images were deposited on SEANOE (DOI: 10.17882/111869).

The type and voucher specimens of *I*. *gracilis* sp. nov. and *I*. *elegans* sp. nov. have been deposited in the Marine Biological Museum of the Chinese Academy of Sciences (MBMCAS) at Qingdao, China. The types of *I*. *pseudarmata* sp. nov. are deposited in the Muséum national d’Histoire naturelle in Paris, France.

### DNA extraction and Sanger DNA sequencing

The TIANamp Marine Animal DNA Kit (Tiangen Biotech, Beijing, China) was used to extract total genomic DNA from polyp tissue. Sequences of the mitochondrial genes *mtMutS* and *cox1*, and the nuclear gene *28S rDNA* were obtained through Sanger sequencing of polymerase chain reaction (PCR) products for molecular systematic analysis. To amplify *mtMutS*, *cox1* and *28S*, the primer pairs AnthoCorMSH (Herrera et al., 2010) and Mut-3458R (Sánchez et al., 2003), COI8414-F (5’-CCAGGTAGTATGTTAGGRGA-3’; McFadden, unpublished) and HCO2198 (Folmer et al., 1994), and 28S-Far and 28S-Rar (McFadden and van Ofwegen, 2013) were used, respectively. PCR conditions were: 2 min at 98 °C followed by 35 cycles (98 °C for 15 s, 45–50 °C for 15 s, and 72 °C for 60 s) and a final extension at 72 °C for 2 min. PCR products were sent to TsingKeBiotech (Beijing, China) for bidirectional sequencing using the corresponding forward and reverse primer. All newly obtained sequences were deposited in NCBI Genbank (Table 2).

**Table 2.** The sequences used in this study. Sequences of new species in this study are given in bold. “–” = data unavailable.

| Species | Voucher | Collection number<br>or isolated<br>fragment number | GenBank Accession numbers |  |  |  |
| --- | --- | --- | --- | --- | --- | --- |
|  |  |  | *mtMutS* | *coxI* | *28S rDNA* | UCEs (SRA) |
| *Isidoides elegans* sp. nov. (holotype) | MBM286500 | M5223 | **PV926094** | **PV915943** | **PV919767** | **SAMN50046240** |
| *I. elegans* sp. nov. (paratype) | MBM286501 | M5043 | **PV926093** | **PV915942** | **PV919768** | **SAMN50046241** |
| *I. gracilis* sp. nov. (holotype, form 1) | MBM286877 | Z076 | **PV926096** | **PV915945** | **PV919769** | **SAMN50046242** |
| *I. gracilis* sp. nov. (paratype, form 1) | MBM286878 | N11 | **PV926095** | **PV915944** | – | **SAMN50046243** |
| *I. gracilis* sp. nov. (paratype, form 1) | MBM286879 | RY16049 | **PV926098** | **PV915946** | **PV919766** | **SAMN50046244** |
| *I. gracilis* sp. nov. (paratype, form 2) | MBM286880 | RY16141 | **PV926097** | **PV915947** | **PV919763** | **SAMN50046245** |
| *I. armata* (form 5) | NIWA163516 | MBM286882 | **PV926100** | **PV915949** | **PV919764** | **SAMN50046247** |
| *I. armata* (form 4) | NIWA82680 | MBM286881 | **PV926099** | **PV915948** | **PV919765** | **SAMN50046246** |
| *I. armata* (form 3) | MNHN-IK-2008-1510 | TER20516 | JN227946,<br>**PV926086** | JN227951,<br>KX530099,<br>**PV915935** | – | **SAMN50046232** |
| *I. armata* (form 3) | AM:G16995 | NORF36/10 | JN228002 | – | – | – |
| *I. armata* (form 3) | MNHN-IK-2009-1632 | CP38321 | KX362340,<br>**PV926090** | **PV915939** | **PV919757** | **SAMN50046236** |
| *I. armata* (form 3) | MNHN-IK-2008-1505 | TER105-1 | JN227934,<br>**PV926081** | **PV915930** | **PV919752** | **SAMN50046227** |
| *I. armata* | – | – | NC_062015 | NC_062015 | – | – |
| *I. armata* | – | – | OL616240 | OL616240 | – | – |
| *I. armata* (form 3) | MNHN-IK-2008-1506 | – | **PV926082** | **PV915931** | **PV919753** | **SAMN50046228** |
| *I. armata* (form 3) | MNHN-IK-2016-2540 | – | **PV926092** | **PV915941** | **PV919762** | **SAMN50046239** |
| *I. armata* (form 3) | MNHN-IK-2016-2539 | – | **PV926091** | **PV915940** | **PV919761** | **SAMN50046238** |
| *I. armata* (form 3) | MNHN-IK-2008-1411 | NOR220652 | **PV926079** | **PV915928** | **PV919750** | **SAMN50046225** |
| *I. armata* (form 3) | MNHN-IK-2008-1504 | TER1012 | **PV926080** | **PV915929** | **PV919751** | **SAMN50046226** |
| *I. armata* (form 3) | MNHN-IK-2008-1507 | TER10084 | **PV926083** | **PV915932** | **PV919754** | **SAMN50046229** |
| *I. armata* (form 3) | MNHN-IK-2011-1000 | – | – | – | – | **SAMN50046237** |
| *I. pseudarmata* sp. nov. (paratype, “*I. armata*” in Pante et al. 2013) | MNHN-IK-2008-1875 | TER1021 | KX362253,<br>**PV926089** | **PV915938** | **PV919760** | **SAMN50046235** |
| *I. pseudarmata* sp. nov. (paratype, “*I. armata*” in Pante et al. 2013) | MNHN-IK-2008-1867 | TER1061 | JX126866,<br>**PV926087** | **PV915936** | **PV919758** | **SAMN50046233** |
| *I. pseudarmata* sp. nov. (paratype) | MNHN-IK-2008-1508 | – | **PV926084** | **PV915933** | **PV919755** | **SAMN50046230** |
| *I. pseudarmata* sp. nov. (paratype) | MNHN-IK-2008-1509 | – | **PV926085** | **PV915934** | **PV919756** | **SAMN50046231** |
| *I. pseudarmata* sp. nov. (holotype) | MNHN-IK-2008-1869 | – | **PV926088** | **PV915937** | **PV919759** | **SAMN50046234** |
| *Parajasonis flabellata* | MBM287336 | M1003 | PP999495 | PP991392 | PP992072 | – |
| *Parajasonis flabellata* | MBM287337 | M1155 | PP999496 | PP991393 | PP992073 | – |
| *Dentatisis bathyalis* | MBM287339 | M088 | PP999498 | PP991395 | PP992075 | – |
| *Dentatisis projecta* | RSIOOC0210 | JL171-B10 | PP999497 | PP991394 | PP992074 | – |
| *Aethisis turriformis* | RSIOOC0241 | JL179-B05 | PP999499 | PP991396 | PP992076 | – |
| *Jasonis thresheri* | TMAK K3879 | OCT052 | JN620806 | JN620807 | – | – |
| *Dokidisis krishanani* | NCPOR/HYD-CIOR/0070 | – | OQ594413 | OQ605881 | OQ625531 | – |
| *Cladarisis nouvianae* | YPM IZ 070870 | RUM1072 | KX362252 | KX530100 | – | – |
| *Keratoisis grayi* | – | 5663 | KX362308 | KX530101 | – | – |
| *Adinisis oblonga* | NCPOR/HYD-CIR/0086 | – | OQ594412 | OQ605880 | OQ625532 | – |
| Keratoisididae sp. S1 | MBM287343 | Z5064 | PQ268061 | PQ248516 | PQ249238 | – |
| Keratoisididae sp. I1 | MBM287345 | YM042 | PQ268063 | PQ248518 | PQ249240 | – |
| Keratoisididae sp. I4 | MBM287348 | YM213 | PQ268066 | PQ248521 | PQ249243 | – |
| Keratoisididae sp. D2 | MBM287352 | M8088 | PQ268070 | PQ248525 | PQ249247 | – |
| Keratoisididae sp. C1 | MBM288057 | 10-4 | PQ268076 | PQ248531 | PQ249253 | – |
| Keratoisididae sp. H1 | MBM288065 | 10-12 | PQ268085 | PQ248540 | PQ249262 | – |
| *Chrysogorgia averta* | – | LII-10-609 | KC788265 | KC788235 | KC788258 | – |
| *Chrysogorgia arboriformis* | MBM286465 | – | ON221842 | ON207094 | ON207672 | – |
| *Parachrysogorgia chrysei* | MBM286358 | – | ON221840 | ON207090 | ON207678 | – |
| *Radicipes stonei* | USNM:IZ:1418<br>007/SW0714 | – | MG986912 | MG986961 | MG980134 | – |

### Genetic distance and phylogenetic analyses of Sanger sequencing

Additional *mtMutS*, *cox1* and *28S rDNA* sequences of the genus *Isidoides*, Keratoisididae and Chrysogorgiidae were obtained from Genbank (Table 2). MAFFT v.7 (Katoh and Standley, 2013) was used with the G-INS-i algorithm to align the sequences and BioEdit v7.0.5 (Hall, 1999) was used to refine the nucleotide alignments (deposited on SEANOE). Genetic distances, calculated as uncorrected “p” distances within- and among species, were estimated using MEGA 11 (Tamura et al., 2021).

Bayesian Inference (BI) and Maximum Likelihood (ML) were used to generate phylogenetic trees of the *mtMutS-cox1* and *28S rDNA* genes. Model selection, Maximum likelihood (ML) and Bayesian inference (BI) analyses were conducted by PhyloSuite v1.2.2 (Zhang et al., 2020; Xiang et al., 2023). The best partitioning scheme and evolutionary models for *mtMuts* and *cox1* were selected for ML using PartitionFinder2 v2.1.1 (Lanfear et al., 2017), with the greedy algorithm and AICc criterion. ModelFinder v2.2.0 (Kalyaanamoorthy et al., 2017) was used to select the best-fit model for BI using the BIC criterion. HKY+F+G4 was selected as the best-fit substitution model for *28S rDNA* alignments. Bayesian Inference (BI) phylogenies were inferred using MrBayes 3.2.7a (Ronquist et al., 2012) using two parallel runs of 10^7^ generations, with 25% of sampled generations discarded as burn-in. Convergence of the MCMC runs was evaluated based on summary statistics in PhyloSuite v1.2.2. Maximum likelihood (ML) phylogenies were inferred using IQ-TREE v3.0.1 (Nguyen et al., 2015) with 1,000 standard bootstrap replicates. Following Hillis and Bull (1993), the ML bootstraps <70%, 70–94% and ≥95% were considered low, moderate and high, respectively. Following Alfaro et al. (2003), the Bayesian posterior probabilities <0.95 and ≥0.95 were considered low and high, respectively.

### DNA Target Enrichment and Sequencing

All twenty-three specimens of *Isidoides* were selected for DNA sequencing of UCEs using target bait enrichment with the ‘octocoral-v2’ baitset, as per Erickson et al. (2021). DNA extracts were sent to Bluescape Hebei Biotech Co. Ltd. (Baoding P.R. China) for target enrichment and library preparation. The library was sequenced by Novogene Co., Ltd. (Beijing) on an Illumina Novaseq X Plus platform; raw sequence reads were deposited in the NCBI Sequence Read Archive (SRA) (https://www.ncbi.nlm.nih.gov/sra) as BioProject PRJNA1293818 and SRA accessions SRX29846129 to SRX29846151. The resulting fastq files were processed using *phyluce* v1.7.3 (Faircloth, 2015) according to Quattrini et al. (2018), except that contigs were assembled with *SPAdes* v.4.0.0 (Bankevich et al., 2012) using the ‘single-cell’ assembly mode without the ‘careful’ option. After matching assembled contigs to the ‘octocoral-v2’ UCE bait set (Erickson et al., 2021), sequence data from three keratoisidid outgroup specimens were obtained from previous studies (McFadden et al., 2022; A. Quattrini pers. comm.) and were included in subsequent alignments and phylogenetic analyses.

Final concatenated alignments (deposited on SEANOE) for phylogenetic inference included loci with 75% and 90% taxon occupancy (i.e., the proportion of taxa represented per locus, hereon called “occupancy matrices”). Each alignment was analyzed independently using BI and ML. For the former, *ExaBayes* v1.5.1 (Aberer et al., 2014) was used with partitioning by locus, using four chains of 10^6^ generations sampled every 1,000 generations. Phylogenetic reconstruction used 90% of the resulting samples, discarding the first 10% as burn-in For the latter, IQ-TREE v3.0.1 (Nguyen et al., 2015) was run on the same locus-based partition scheme as ExaBayes, using Model Finder Plus (MFP+MERGE) for model selection, 1000 ultrafast bootstrap replicates with --bnni option to reduce the risk of overestimating branch supports, and 1000 SH-aLRT tests.

### Species delimitation

Species delimitation was conducted using an integrative framework combining phylogenomic inference under the multispecies coalescent (ASTRAL+SODA), alongside multivariate (PCA) and model-based (STRUCTURE) analyses of single nucleotide polymorphisms (SNPs). First, IQ-TREE was used to produce one tree per UCE locus, using MFP+MERGE as above. These gene trees where then used as input in ASTRAL v5.7.8 (Zhang et al., 2018) to infer a species tree with branch lengths computed as coalescent units (Sayyari and Mirarab, 2016), which in turn was used as a guide tree for SODA v1.0.2 (Rabiee and Mirarab, 2021). SODA tests the null hypothesis that each internal branch has length zero, i.e., that internal branches corresponds to a polytomy under the multispecies coalescent. The significance level α determines the threshold for rejecting this null hypothesis. We set α to 0.05 (default) and 0.01 (stringent). SODA uses the polytomy test developed by Sayyari and Mirarab (2018) and implemented in ASTRAL. Groups were not defined a priori as to not influence the delimitation results. Finally, as the behavior of SODA is known to be affected by the number of loci (Lähteenaro et al., 2024) and, most probably, occupancy, we ran SODA on 75%, 90% and 100% occupancy matrices.

The SNP calling pipeline (archived on SEANOE and https://github.com/ericpante/isidoides) is based on https://github.com/Lavarchus/SNP-calling-GATK4 and Erickson et al. (2021). First, MBM286878 was selected as the reference individual for SNP calling, as it contained the largest number of captured UCEs. Clean reads from the 21 *Isidoides* individuals selected (see above) were mapped to MBM286878 contigs using bwa-mem2 (v2.2.1; Vasimuddin et al., 2019); unmapped reads were removed and mapped reads were sorted using samtools (v1.21; Li et al., 2009). Duplicate reads were removed and the resulting bam files were indexed with picard (v2.23.5; https://broadinstitute.github.io/picard/). gatk4 (v4.6.1.0; McKenna et al., 2010) was used to perform the core SNP calling, namely individual-based haplotype calling (HaplotypeCaller function), consolidation (CombineGVCFs), joint genotyping of all individuals (GenotypeGVCFs) and SNPs and indels extraction (SelectVariants). Variants were hardfiltered (VariantFiltration) according to recommendations from the Broad Institute (QD < 4 for SNPs, QD < 2 for indels, SOR>10, QUAL < 30.0, MQ < 40). The BQSR (base quality score recalibration) procedure was applied to correct for sequencing errors (BaseRecalibrator, ApplyBQSR) and the adjusted quality scores were compared to original scores with AnalyzeCovariates and the pipeline (from the HaplotypeCaller step to the VariantFiltration step) was re-run using corrected base quality scores. Two repetitions of the pipeline were sufficient to reach convergence of pre- and post-recalibration quality covariates. One dataset was exported without filtering for allele frequencies, another with a minor allele frequency cutoff of 5%. The resulting datasets were filtered to retain one biallelic SNP (minimum distance of 1500 nt between variants so most SNPs come from a single UCE locus) with no missing data (vcftools v0.1.16; Danecek et al., 2011).

plink (v1.90b6.18, Shaun Purcell & Christopher Chang, www.cog-genomics.org/plink/1.9/; Chang et al., 2015) was used to export the SNP data in the Structure format and to perform the PCA; plotting was done in R (v4.5.2; R Core Team, 2025). Structure (v2.3.4; Pritchard et al., 2010) assigns individuals into K groups based on Hardy-Weinberg equilibrium and linkage disequilibrium. Its admixture model was used here to delineate species-level units by identifying genetically cohesive clusters consistent with interbreeding populations (Singhal et al., 2025). Twenty replicate runs were used for each value of K (1 to 8; see below). Each run consisted of 400,000 generations, including a burn-in of 200,000 generations.

Introgression among species was tested using the SNP dataset and gene trees computed with iq-tree. Dsuite (v0.5_r53; Malinsky et al., 2020) was used to compute the D-statistic based on the final VCF file described above (no MAF filter). Ghostparser (v2026-04-01; Tolman and Suvorov, 2025) was used to detect introgression among sampled species, as well as "ghost introgression" between sampled and unsampled (or extinct) taxa. The Ghostparser method is based on a combination of tree topology and branch lengths, under the multi-species coalescent framework. Gene trees generated from the 75% occupancy matrix were used to maximise the number of triplets and the species tree was based on the 100% occupancy matrix (ML analysis detailed above). For both analyses, p-values were adjusted with the FDR method implemented in R (stats::p.adjust) and interpreted at α=0.01 to avoid false positives.

## Results

### Genetic distance and phylogenetic analyses of Sanger sequencing

The new sequences, including 22 *mtMutS*, 22 *cox1* and 20 *28S rDNA*, were obtained successfully from 23 specimens and deposited in GenBank (Table 2). The alignments comprised 808, 540 and 682 bp nucleotide positions for the aforementioned regions, respectively. Based on these alignments, the *mtMutS*-*cox1* and *28S* rDNA interspecific distances among the *Isidoides* species ranged from 0–0.34% and 0–3.46%, respectively, and the corresponding intraspecific distances were 0–0.17% and 0–0.47% (Tables 3, 4).

**Table 3.** Inter- and intraspecific uncorrected pairwise (p) distances among Isidoidae species at *mtMutS-cox1* (%)

| Species | 1 | 2 | 3 | 4 | 5 |
| --- | --- | --- | --- | --- | --- |
| 1 *Isidoides armata* (form 3: MNHN-IK-2008-1504, 2008-1505, 2008-1506, 2009-1632, 2016-2539, NORF3610, TER1051, CP38321; form 4: NIWA82680; NC062015, OL616240; form 5: NIWA163516), *Isidoides pseudarmata* sp. nov. (MNHN-IK-2008-1508, 2008-1509, 2008-1867, 2008-1869, 2008-1875, TER1021, TER1061) | 0 |  |  |  |  |
| 2 *Isidoides armata* (form 3: MNHN-IK-2008-1411, 2008-1507, 2008-1510, 2016-2540) | 0.08 | 0 |  |  |  |
| 3 *Isidoides gracilis* (form 1: Z076, N11, RY16049; form 2: RY16141) | 0.08 | 0.17 | 0 |  |  |
| 4 *Isidoides elegans* M5043, M5223 | 0.17 | 0.25 | 0.08 | 0 |  |
| 5 *Isidoides armata* (form 3: TER20516) | 0.17 | 0.08 | 0.25 | 0.34 | - |

**Table 4.** Inter- and intraspecific uncorrected pairwise (p) distances among Isidoidae species at *28S rDNA* (%).

| Species | 1 | 2 | 3 | 4 | 5 | 6 | 7 | 8 | 9 | 10 | 11 |
| --- | --- | --- | --- | --- | --- | --- | --- | --- | --- | --- | --- |
| 1 *Isidoides armata* (form 3: MNHN-IK-2008-1505, 2008-1506; form 5: NIWA163516) | 0 |  |  |  |  |  |  |  |  |  |  |
| 2 *I. armata* (form 3: MNHN-IK-2008-1504) | 0.16 | - |  |  |  |  |  |  |  |  |  |
| 3 *I. armata* (form 3: MNHN-IK-2008-1507, 2016-2540; form 4: NIWA82680) | 0.16 | 0.31 | 0 |  |  |  |  |  |  |  |  |
| 4 *I. armata* (form 3: MNHN-IK-2008-1632, 2016-2539) | 0.31 | 0.16 | 0.16 | 0 |  |  |  |  |  |  |  |
| 5 *I. armata* (form 3: MNHN-IK-2008-1411) | 0.31 | 0.47 | 0.16 | 0.31 | - |  |  |  |  |  |  |
| 6 *I. pseudarmata* sp. nov. MNHN-IK-2008-1509, 2008-1867, 2008-1869 | 0.63 | 0.79 | 0.47 | 0.63 | 0.63 | 0 |  |  |  |  |  |
| 7 *I. pseudarmata* sp. nov. MNHN-IK-2008-1508 | 0.79 | 0.94 | 0.63 | 0.79 | 0.79 | 0.16 | - |  |  |  |  |
| 8 *I. pseudarmata* sp. nov. MNHN-IK-2008-1875 | 0.94 | 1.10 | 0.79 | 0.94 | 0.94 | 0.31 | 0.16 | - |  |  |  |
| 9 *Isidoides gracilis* sp. nov. (form 1: RY16049, Z076; form 2: RY16141) | 2.52 | 2.68 | 2.36 | 2.52 | 2.52 | 2.83 | 2.99 | 2.83 | 0 |  |  |
| 10 *Isidoides elegans* sp. nov. M5223 | 2.83 | 2.99 | 2.68 | 2.83 | 2.83 | 3.15 | 3.31 | 3.15 | 0.94 | - |  |
| 11 *I. elegans* sp. nov. M5043 | 2.99 | 3.15 | 2.83 | 2.99 | 2.99 | 3.31 | 3.46 | 3.31 | 1.10 | 0.16 | - |

The ML trees were nearly identical in topology to the BI trees, and thus a single tree with both support values is presented for concatenated *mtMutS-cox1* and *28S rDNA* (Figs. 2, 3). All the *Isidoides* species formed a monophyletic group in both the *mtMutS-cox1* and *28S rDNA* trees with high to full branch support. In the *mtMutS-cox1* trees (Fig. 2), two specimens of *I. elegans* sp. nov. formed a single clade with high support in the BI tree (BI posterior probability = 0.98), and were sister to *I. gracilis* sp. nov., albeit with low support. Two forms of *I. gracilis* sp. nov. clustered together with low support. *Isidoides pseudarmata* sp. nov. clustered with three forms of *I. armata* species with low support.

**Figure 2.**
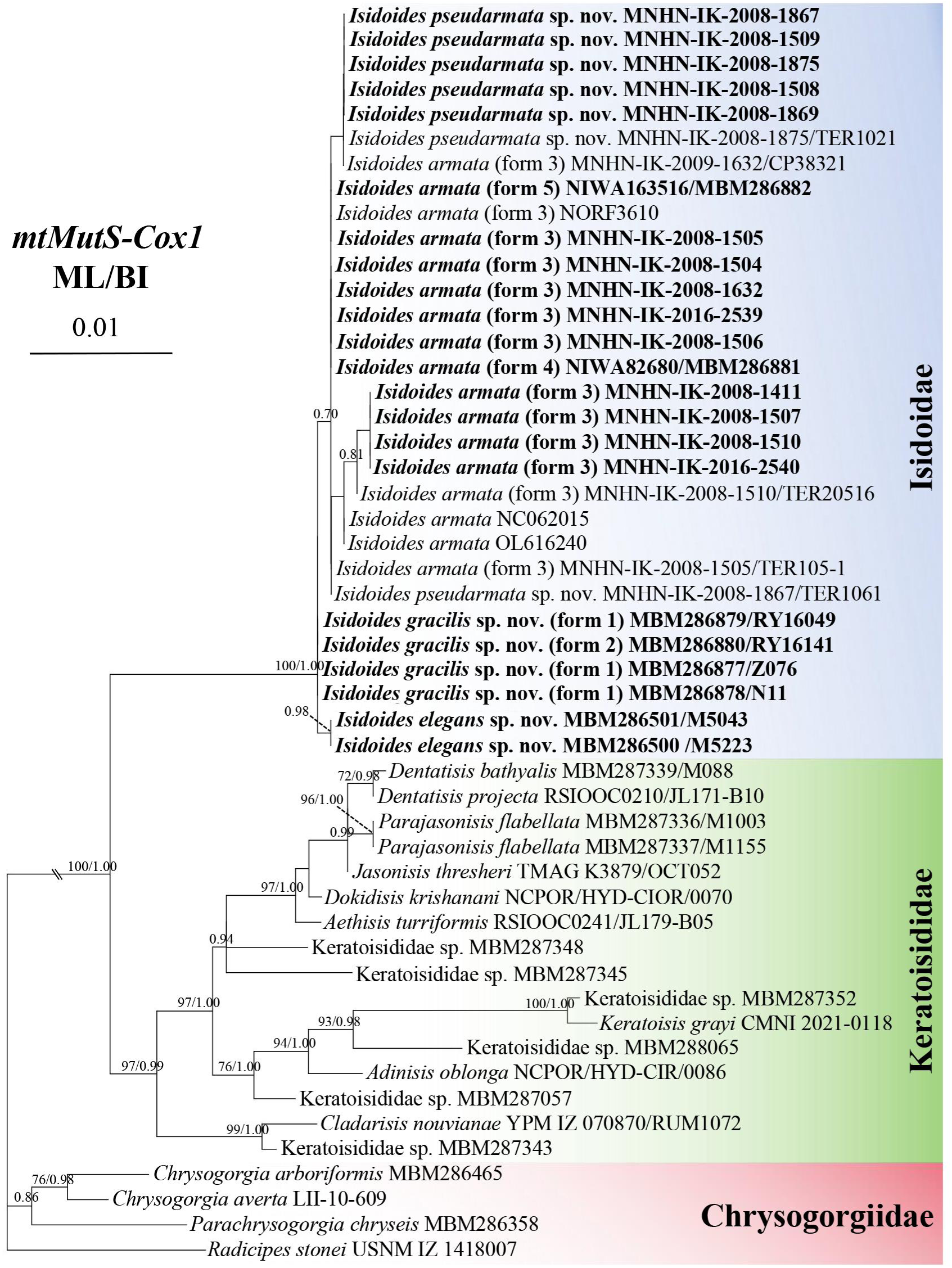
Maximum Likelihood (ML) tree of the *mtMutS-cox1* marker constructed from a 1348 bp alignment, showing phylogenetic relationships among the isidoid, keratoisidid and chrysogorgiid species. The Bayesian Inference (BI) tree is identical to the ML tree in topology. BI posterior probabilities (after slash) and ML bootstrap values (before slash) are shown on each branch. Branches with ML bootstrap <70% or BI posterior probability <0.70 do not display support values. Chrysogorgiidae species are used as the outgroup. Voucher/isolate numbers are listed at the right-hand side of the cladogram. The specimens with sequences generated in this study are presented in bold.

In the *28S rDNA* trees (Fig. 3), two specimens of *I. elegans* sp. nov. formed a single clade with high to full branch support in both the ML and BI trees (BI posterior probability =1.00, ML bootstrap = 100). Three specimens of both forms of *I. gracilis* sp. nov. formed a single clade with moderate to high support in the BI and ML trees (BI = 0.89, ML = 99), and this clade was sister to *I. elegans* sp. nov. Five specimens of *I. pseudarmata* sp. nov. formed a single clade with full and moderate support in the BI and ML trees (BI = 1.00, ML = 91), respectively. Three forms of *I. armata* clustered together with low support. Concatenating the *mtMutS-cox1* and *28S* alignment did not result in additional resolution.

**Figure 3.**
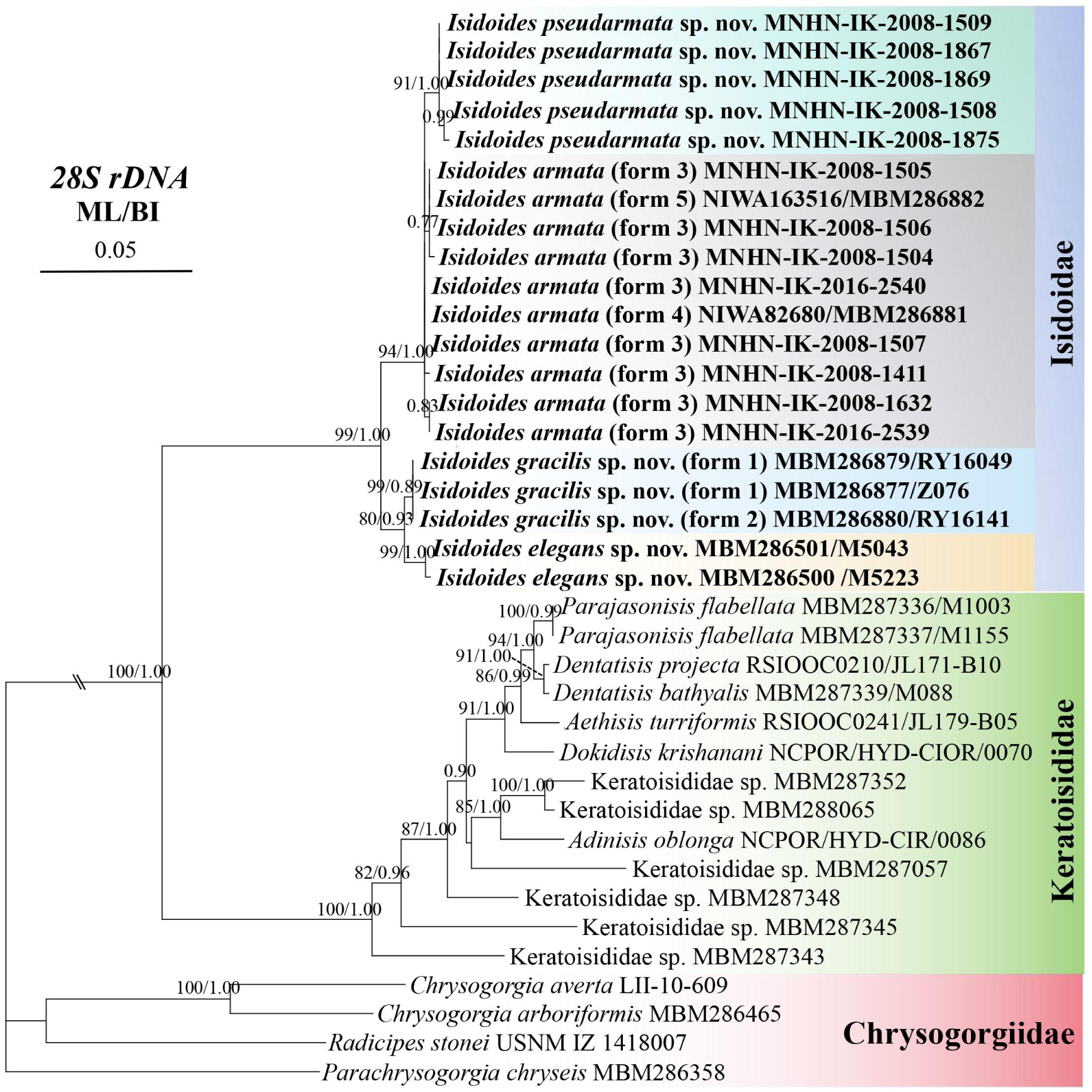
Maximum Likelihood (ML) tree of the *28S* gene constructed from a 682 bp alignment, showing phylogenetic relationships among the isidoid, keratoisidid and chrysogorgiid species. The Bayesian Inference (BI) tree is identical to the ML tree in topology. BI posterior probabilities (after slash) and ML bootstrap values (before slash) are shown on each branch. Branches with ML bootstrap <70% or BI posterior probability <0.70 do not show support values. The out groups are the species of Chrysogorgiidae. Voucher/isolate number are listed at the right side of the cladogram. The specimens with their sequences presented in this study are in bold.

### Phylogenomic analysis of UCEs sequences

After quality-based trimming, the number of reads generated for target-enriched sequences ranged from 5,052,234 to 28,465,519 for each of the 23 *Isidoides* specimens (Supplemental file). Trimmed reads were assembled *de novo* into an average of 125,486 (SE=16,877) contigs per sample, with a range of 32,906 to 295,268. Matching to the ‘octocoral-v2’ baitset resulted in the recovery of 176 to 2,250 UCE loci (x̄=1788, SE=109). Two specimens that each produced fewer than 600 UCE loci (MNHN-IK-2011-1000, 2016-2539, collected in 1985 and 2008, respectively) were discarded from further analysis since their inclusion reduced the resolution and support for resulting phylogenetic trees (data not shown). Sequence data for the remaining 21 specimens were concatenated to produce final alignments of 932,219 bp for 1,572 loci (75% occupancy) and 430,746 bp for 762 loci (90% occupancy). Multi-locus ML and Bayesian phylogenetic inferrences produced topologies that were identical (Fig. 4). ML and Bayesian trees reconstructed from the 75% occupancy matrix alignment produced identical results, except for two unsupported branches (<80%) in *I. armata*, one in *I. pseudarmata* and one in *I. gracilis*; within *I. armata*, MBM286881 was sister to all form 3 instead of grouping with MNHN-IK-2008-1510.

**Figure 4.**
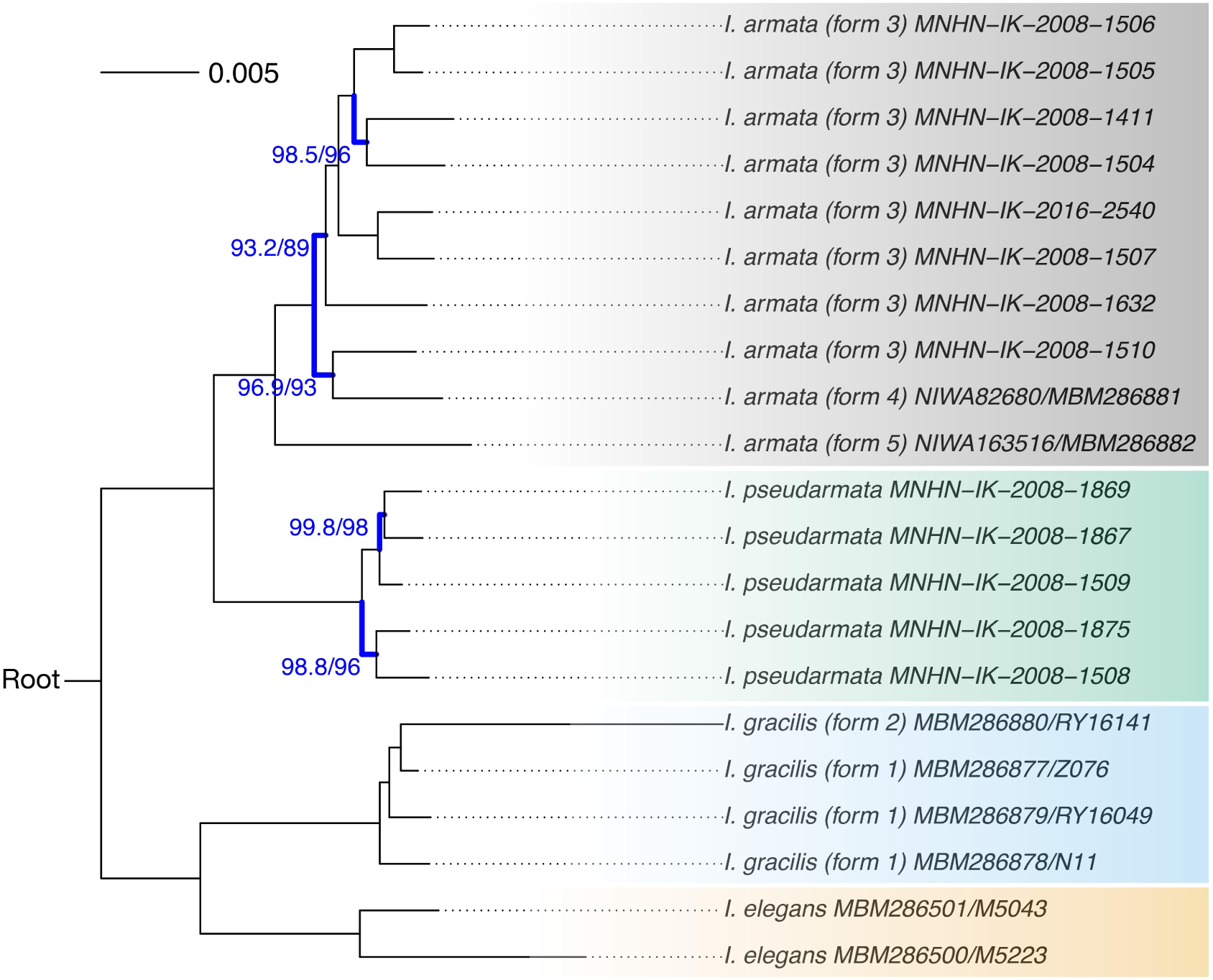
Maximum Likelihood (ML) tree inferred from the 90% occupancy matrix based on UCE loci. Branch support is present with SH-aLRT on the left and ultrafast bootstrap on the right, in percent; only values different from 100% are shown and branches bolded in blue. The bayesian tree topology inferred with Exabayes is identical and all branches had 100% posterior probability support. The tree was rooted using three keratoisidids.

Tree topologies for the UCE analysis distinguished four well-supported clades, corresponding to *I. armata* plus three new species described herein. The UCE analysis resolved *I. gracilis* sp. nov. with two forms, *I. pseudarmata* sp. nov. and *I. elegans* sp. nov. as monophyletic clades, as seen in the *mtMutS-cox1* tree (for *I. elegans* sp. nov.: Fig. 2) and the *28S* tree (for all new species: Fig. 3), and fully resolved the relationships among *I. armata* specimens with three forms, which formed a polytomy in both single-locus gene trees. Further subdivisions within each of these clades were also well-supported but morphological analysis was unable to identify discriminating characters and as such these genetic differences were interpreted as intraspecific variation within the proposed species diagnoses.

### Species delimitation under the multispecies coalescent

The 100% occupancy matrix contained 65 loci and was 37,719 bp long. The ASTRAL species trees recovered from each occupancy matrix were identical, except for some relationships within *I. pseudarmata* sp. nov. and *I. armata* (100% vs 90% and 75%). Within these clades, branch lengths were short (reflecting little time in coalescent units) and poorly supported by local posterior probabilities, which is expected at the intra-specific level. At α=0.05, SODA recovered 14, 12 and eight putative species within *Isidoides*, for the 75, 90 and 100% occupancy matrices, respectively. At α=0.01, SODA recovered 12, 12 and five putative species, for the 75, 90 and 100% occupancy matrices, respectively. As 75% and 90% occupancy matrices led to over-splitting, only results from the 100% occupancy dataset are retained (Fig. 5). In both analyses, *I. elegans* sp. nov., *I. pseudarmata* sp. nov., *I. gracilis* sp. nov. form 1 and *I. gracilis* sp. nov. form 2 were each identified by SODA as putative species. Analyses at α=0.01 and α=0.05 differed in the delimitation of some *I. armata* specimens, the former lumping them together as one species, and the latter splitting *I. armata* form 5, *I. armata* form 4, and two clades of form 3 (MNHN-IK-2008-1507 and MNHN-IK-2016-2540 on one hand, all other form 3 in the other; Fig. 5).

**Figure 5.**
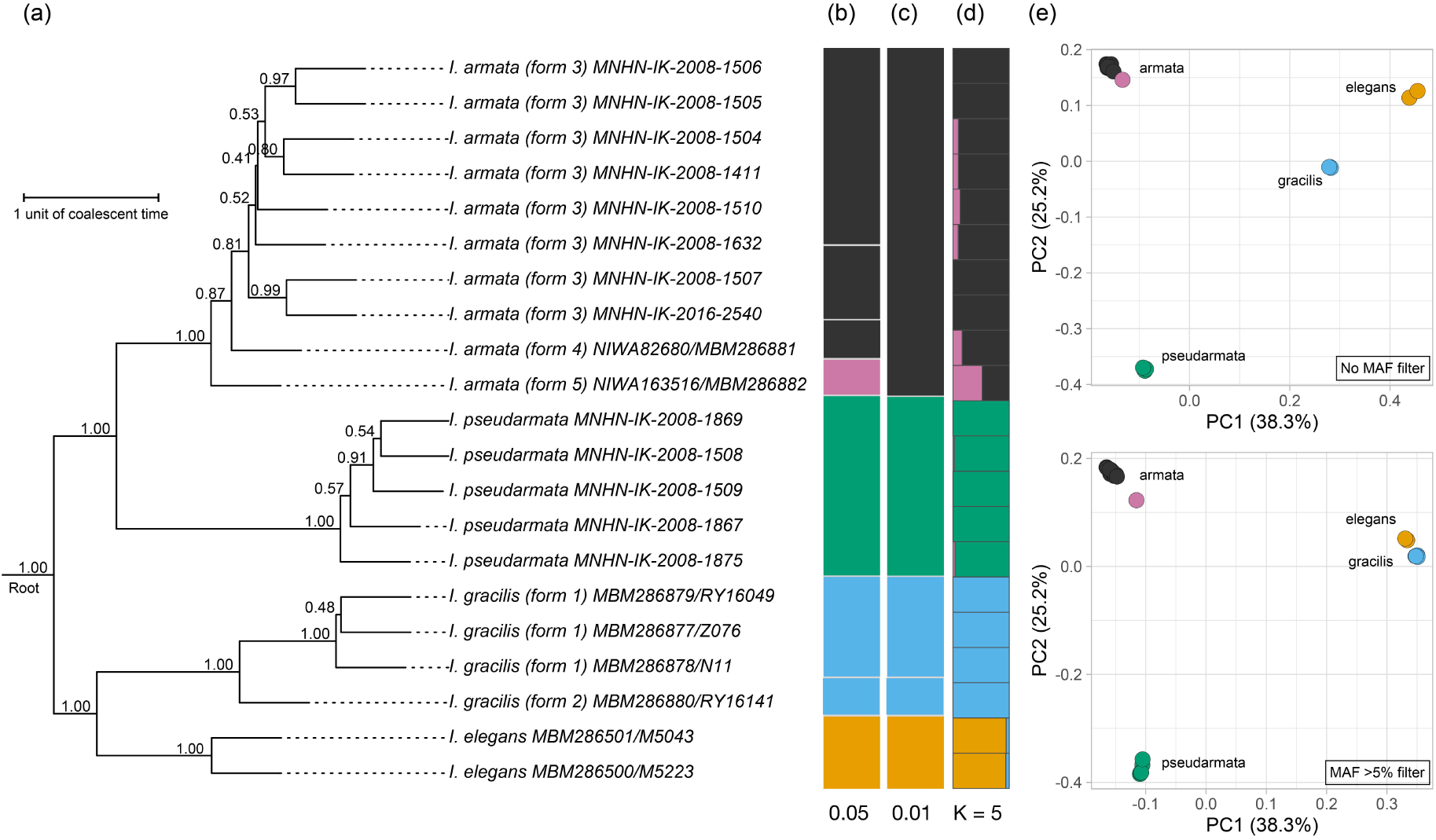
Integrative genetic species delimitation. (a) ASTRAL species tree based on the 100% occupancy matrix, rooted to keratoisidids. Branch support represents local posterior probabilities and internal branch lengths are in coalescent units (terminal branches plotted with arbitrary length). (b-c) SODA delimitations at α=0.01 and α=0.05, respectively (100% occupancy matrix). (d) Structure plot (admixture model) at K=5. (e) PCA on the no-MAF (top) and MAF (bottom) SNP datasets. Numbers in parentheses correspond to the percentage of variance explained by each axis.

A total of 2021 unlinked, biallelic SNPs were recovered (no MAF dataset). When variants were filtered for a MAF>5%, 2089 SNPs were retained (MAF dataset). Based on the no MAF dataset, specimens cluster according to the four putative species (Fig. 5). PC1 separated taxa from the *pseudarmata*/*armata* and *elegans*/*gracilis* clade, and PC2 separated individual putative species; no further information was found in PC3. The same result was observed based on the MAF dataset, although *elegans* and *gracilis* cluster closer to each other, while remaining discrete units. Structure runs on the no MAF data suggest the presence of four clusters corresponding to our four putative species (Fig. 5 for K=5; Supplementary file for K=2 to K=8 for both datasets). For K>5, no additional cluster was formed. Starting at K=5, *I. armata* form 5 (MBM286882) was characterised by 65% cluster *armata*, while the remaining 35% were private to this individual, suggesting that its is a hybrid between *I. armata* and an unsampled taxon (Supplemental file). This pattern does not hold when the analysis is run on the MAF dataset, most *I. armata* individuals being characterized by two clusters, the less prevalent cluster being more prevalent in forms 4 and 5, compared to form 3 (Fig. 5). This pattern holds at higher values of K (Supplemental file).

Weak evidence consistent with introgression within the *armata*/*pseudarmata* clade was detected using the D-statistic (Z>2.5, p<0.01) on the SNP dataset (no MAF), but this signal was not significant after FDR correction (adjusted p=0.056). After filtering GhostParser results to retain only triplets consistent with the species tree, significant Discordant Count Tests (DCT) were recovered mainly for triplets in which *elegans* was the outgroup to the *armata-pseudarmata* pair, suggesting excess allele sharing between *elegans* and either *pseudarmata* or *armata*. GhostParser also revealed signals consistent with unsampled introgression (significant Tree Height Test, THT), particularly involving *I. gracilis* individuals.

### Systematics

**Taxonomy**

**Phylum Cnidaria Hatschek, 1888**

**Subphylum Anthozoa Ehrenberg, 1834**

**Class Octocorallia Haeckel, 1866**

**Order Scleralcyonacea McFadden, van Ofwegen & Quattrini, 2022**

**Family Isidoidae Heestand Saucier, France & Watling, 2021**

### Genus *Isidoides* Nutting, 1910

#### Diagnosis (Based on Heestand Saucier et al., 2021)

Colony planar with pseudo-dichotomous branching, sometimes covered with a layer of thick tegument. Axis solid and calcareous, white to dark golden-brown, not articulated, occasionally flattened along the face of the fan. Polyps non-retractile, large tubular or small wart-like, some of them distally expanded, biserially or irregularly arranged on all sides. Sclerites as elongate scales, flattened rods and rods with relatively uniform shape, occasionally cross-shaped. Sclerites abundant and tightly packed on the polyp and coenenchyme, but absent from the pharynx.

#### Type species

*Isidoides armata* Nutting, 1910.

#### Distribution

Indo-West Pacific and southwest Pacific, 424–1065 m (Table 1; Nutting, 1910, Pante et al., 2013, and present study).

### Isidoides armata Nutting, 1910

Fig. 6D, E, G, N; 7A–C; 8–14

**Figure 6.**
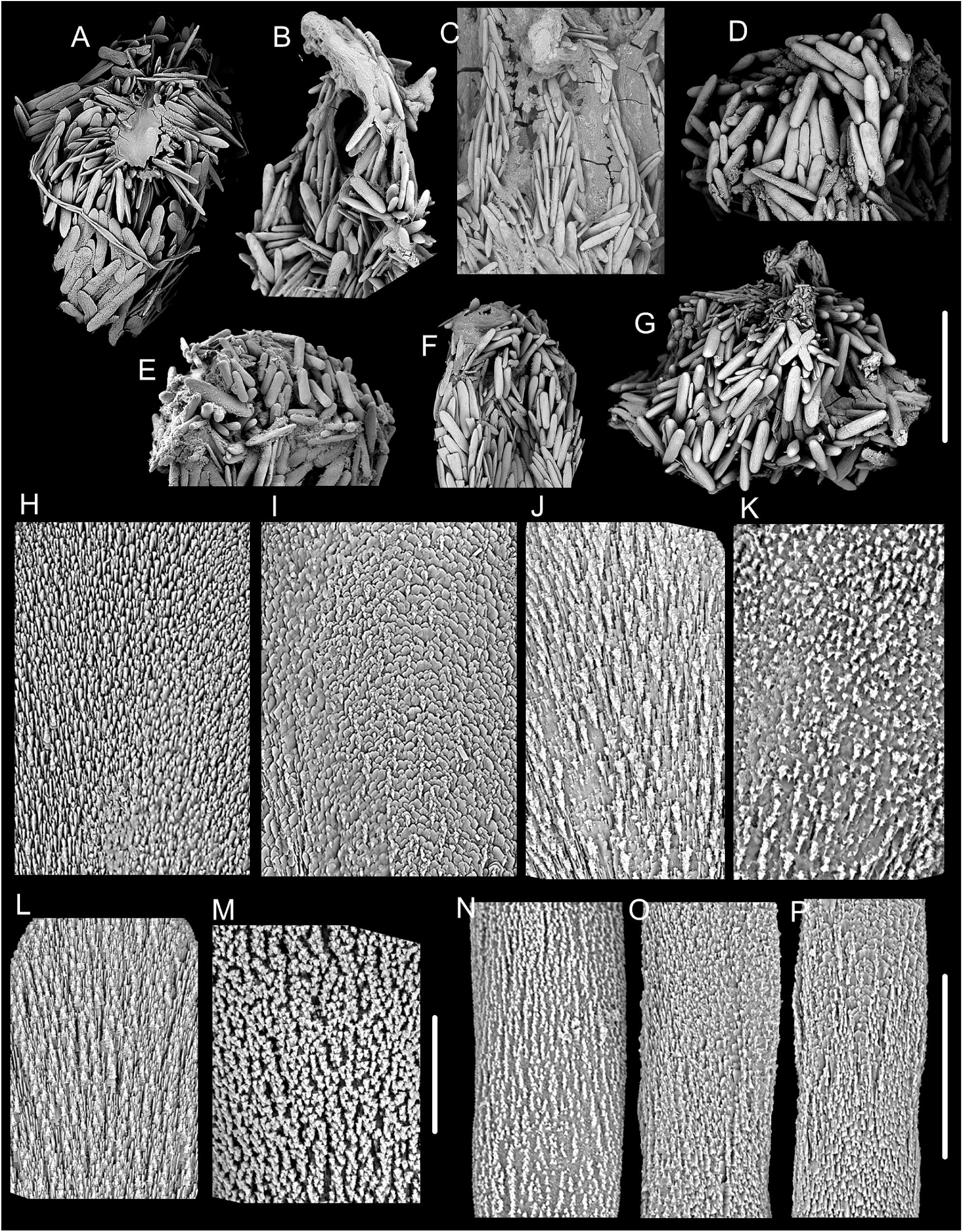
Morphology of tentacular sclerites. **(A**–**G), surfaces of the sclerites in the polyp body wall and coenenchyme (H**–**M) and in tentacles (N**–P) of different *Isidoides* species. (A) flattened rods in *I. elegans* sp. nov.; (B, C) flattened rods in *I. gracilis* sp. nov. (specimen MBM286877 and MBM286880, respectively); (D) rods in *I. armata* (specimen NIWA82680); (E) rods in *I. armata* (paralectotype); (F) rods in *I. pseudarmata* sp. nov. (MNHN-IK-2008-1509); (G) rods in *I. armata* (NIWA163516); (H) striped; (I) lamellar; (J) feather-shaped; (K) granulated; (L) mixed shaped; (M) densely granulated; (N) ridged and granulated in *I. armata*; (O, P) non-ridged and non-granulated in *I. gracilis* sp. nov. and *I. elegans* sp. nov., respectively, as in *I. pseudarmata* sp. nov. Scale bar: 500 μm (A–G), 50 μm (N–P), 30 μm (H–M).

**Figure 7.**
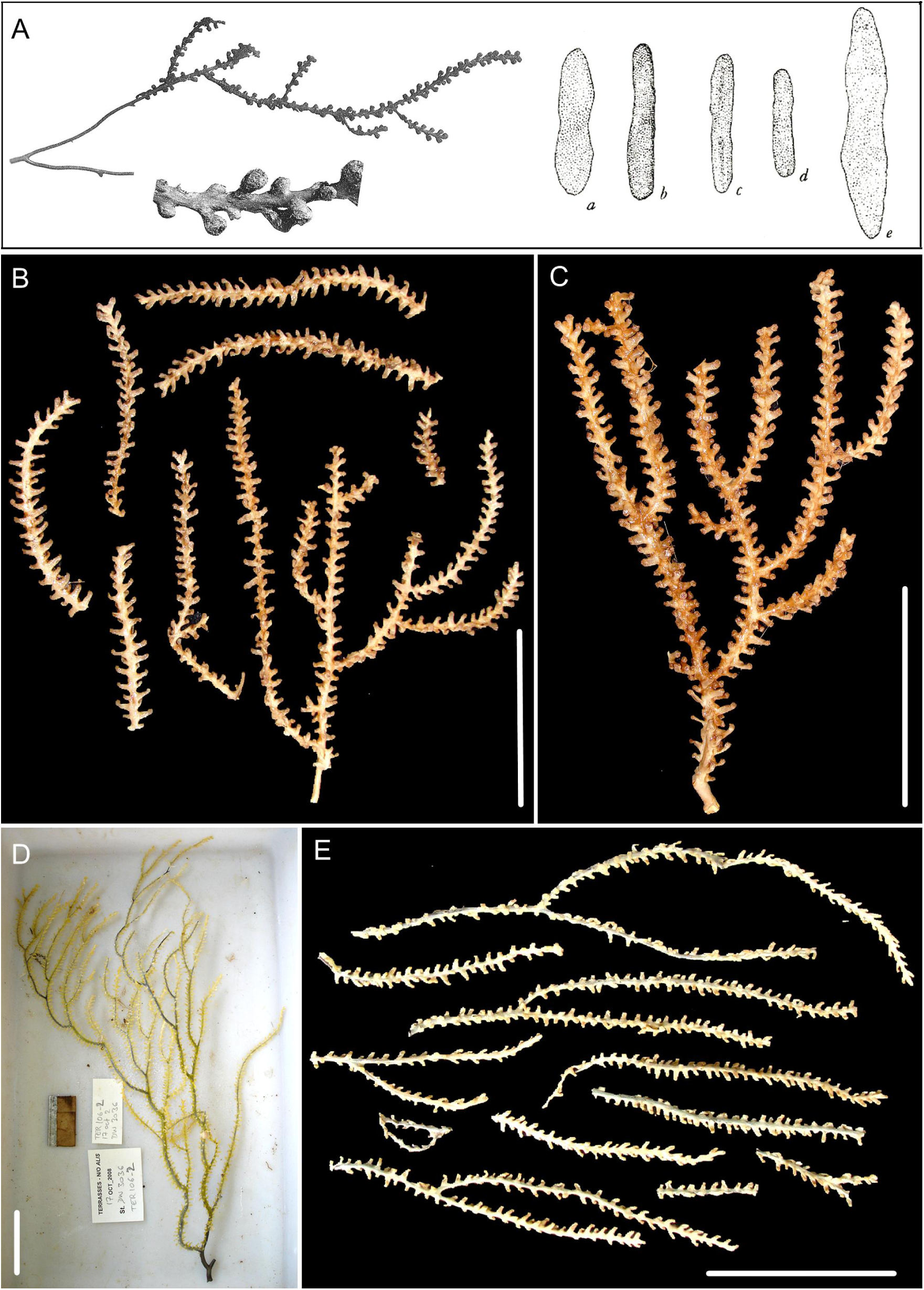
The lectotype of *Isidoides armata* Nutting, 1910 and some recent collections identified as “*I. armata*” by Pante et al. (2013). (A) Nutting’s original illustration of *I. armata* with original figure numbering, including a large branch (15 cm long and 5 cm wide), a part of the branch with close-up of polyps and sclerites (a–e, with e being 454.5 μm tall); (B) specimen MNHN-IK-2008–1505 (TER1051) after fixation, figure processed from Fig. 4D in Pante et al. (2013); (C) specimen MNHN-IK-2008-1510 (TER20516) after fixation, figure processed from Fig. 4E in Pante et al. (2013); (D) specimen TER1062 after collection, figure processed from Fig. 5A in Pante et al. (2013); (E) specimen MNHN-IK-2008-1869 after fixation. Scale bar: 5 cm (B–E).

*Isidoides armata* Nutting, 1910: 33–34, Plate VIII, figs. 2, 2a, Plate XI, fig. 5.

*Isidoides armata*: Pante et al., 2013: 371–374, figs. 2–4, 8–10; Heestand Saucier et al., 2021: 268 (listed only); McFadden et al., 2022: 8, 11 (listed only).

**Figure 8.**
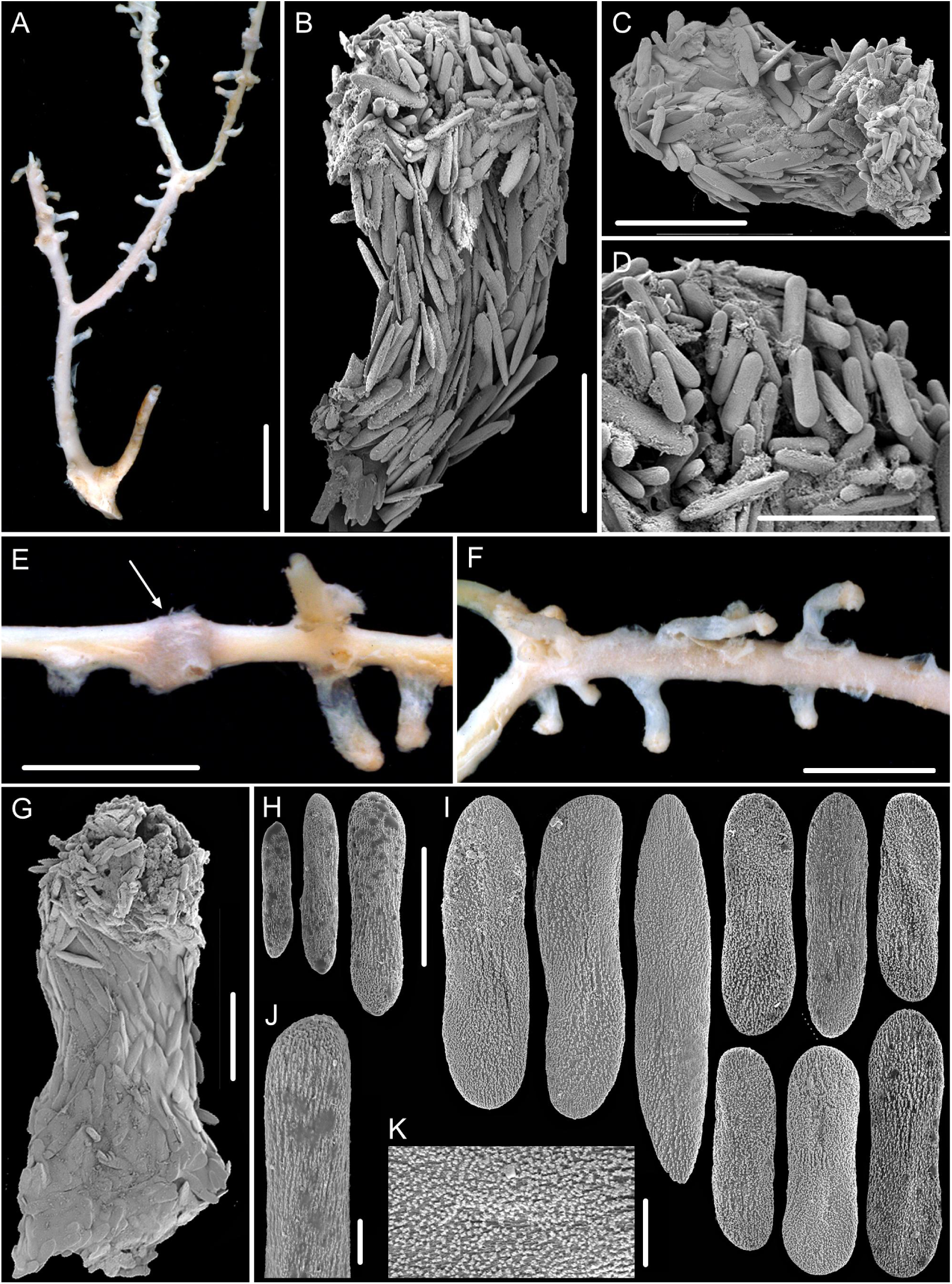
External morphology, polyps and sclerites of the paralectotype of *I. armata* form 2 (specimen ZMA Coel. 02711). To protect the remaining paralectotype specimens, only a few polyps and sclerites were selected for imaging. (A) damaged fragment with few remaining polyps, with scars on the branch coenenchyme indicating that many polyps have fallen off the colony (Pante et al. 2013); (B, C, G) single polyps under SEM, among them, (B) figure processed from Fig. 3C in Pante et al. (2013); (E, F) part of a branch with a few polyps under a light microscope, with a feature resembling a proteinaceous node (arrow) in E, which was processed from Fig. 3B in Pante et al. (2013); (H) small rods in tentacles under SEM; (I) flattened rods in polyp body wall and coenenchyme under SEM; (J, K) close-up of the surface of tentacle rods and polyp flattened rods under SEM, respectively. Scale bar: 1 cm (A), 5 mm (E, F), 500 μm (B, C, G), 300 μm (D), 100 μm (H and I in the same scale), 20 μm (J, K).

**Figure 9.**
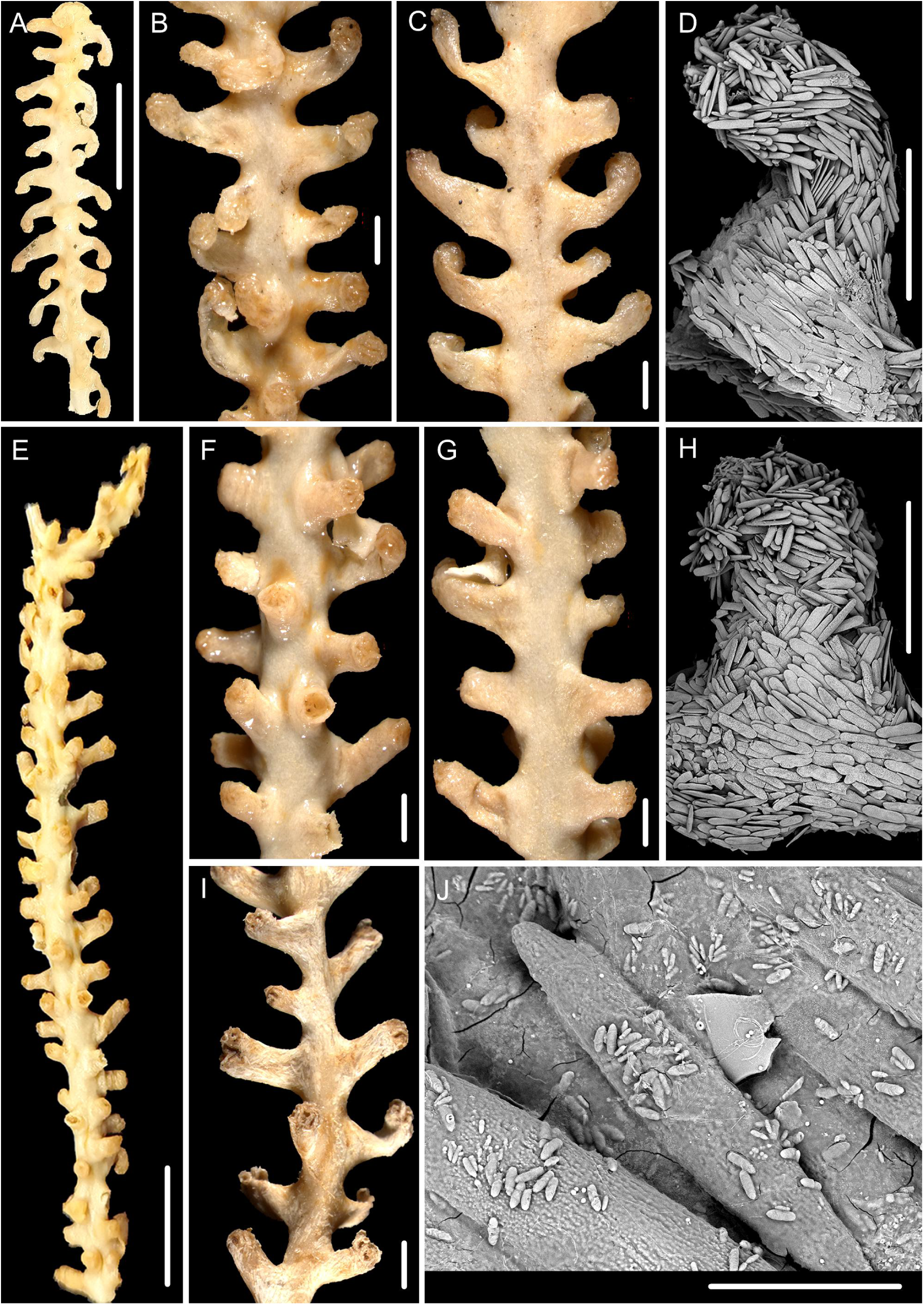
The external morphology and polyps of *I. armata* form 3, specimen MNHN-IK-2008-1505. **(A**–**D), 2009-1632 (E**–**H) and 2008-1411 (I, J).** (A, E) separated fragment; (B, F, I) the front side of branch under a light microscope; (C, G) the back side of branch under a light microscope; (D, H) single polyp under SEM; (J) nematocysts under SEM. Scale bars: 1 cm (A, E); 2 mm (B, C, F, G, I); 1 mm (D, H), 100 μm (J).

**Figure 10.**
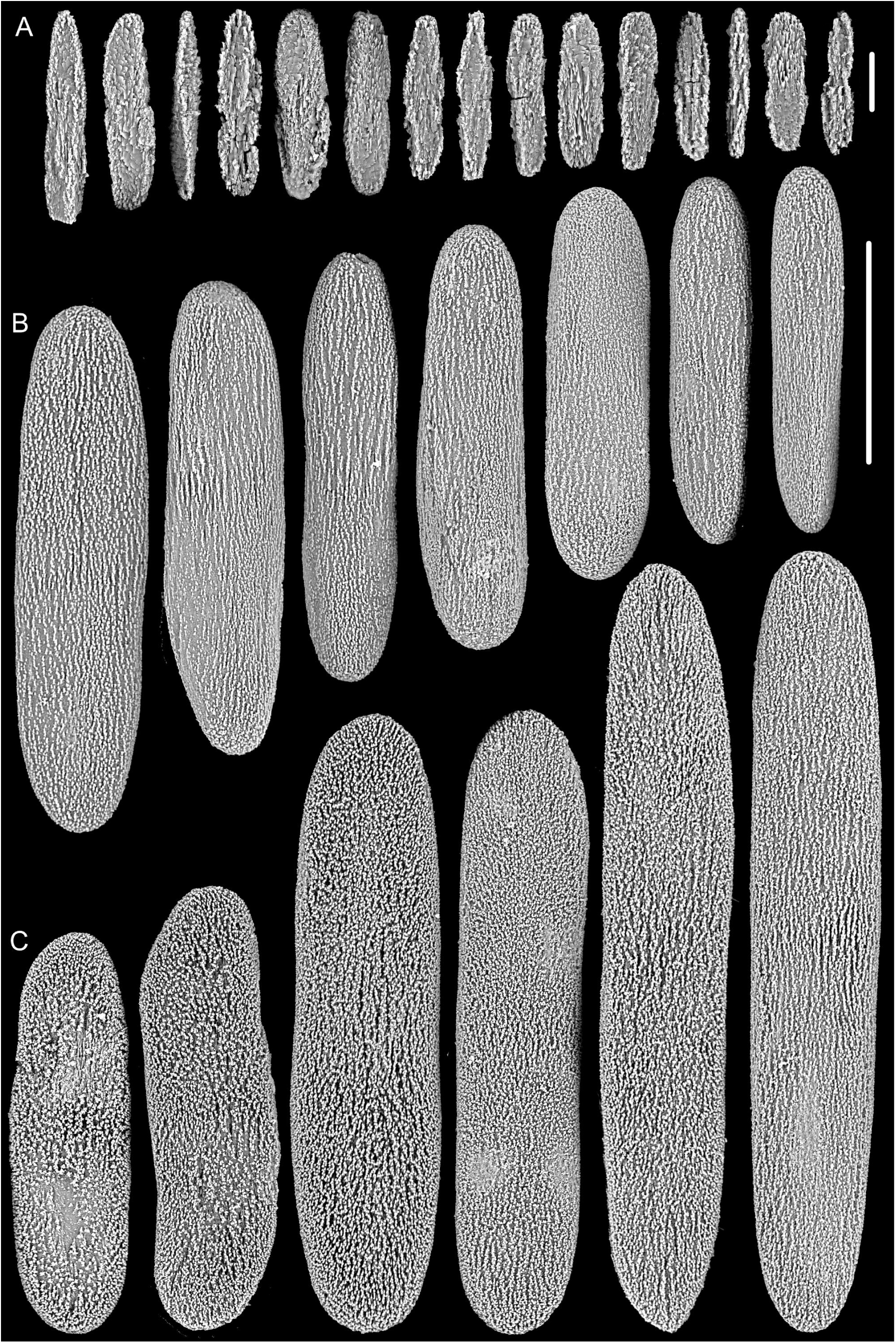
Sclerites of *I. armata* form 3 (specimen MNHN-IK-2009-1632). (A) small flattened rods in tentacles; (B) large rods in tentacles; (C, D) flattened rods in the polyp body wall and coenenchyme. Scale bar: 100 μm (B and C in the same scale), 20 μm (A).

#### Material examined: Paralectotype

ZMA Coel. 02711, station 297 (10.650°S, 123.670°E), south of the western tip of Timor, 520 m. Other specimens: MNHN-IK-2008-1505, station DW3035 (22.687°S, 168.942°E), Walpole Island, Loyalty Ridge, New Caledonia, 790–800 m, 16 October 2008. MNHN-IK-2008-1411, station DW2065 (25.267°S, 168.933°E), Athos Seamount, Norfolk Ridge, New Caledonia, 750–800 m, 26 October 2003. MNHN-IK-2008-1504, station DW3031 (22.675°S, 168.962°E), Walpole Island, Loyalty Ridge, New Caledonia, 720 m, 16 October 2008. MNHN-IK-2009-1632, station CP3832 (21.993°S, 167.120°E), south-eastern slope of New Caledonia, 645 m, 8 September 2011. MNHN-IK-2008-1507, station DW3097 (21.996°S, 167.105°E), south-eastern slope of New Caledonia, 500–941 m, 16 October 2008. MNHN-IK-2008-1506, station DW3035 (22.687°S, 168.942°E), Loyalty Ridge, New Caledonia, 790–800 m, 16 October 2008. MNHN-IK-2008-1510, station DW3041 (23.958°S, 169.722°E), Mont J, Loyalty Ridge, New Caledonia, 840–800 m, 17 October 2008. MNHN-IK-2016-2540, station DW3097 (22.000°S, 167.100°E), south-eastern slope of New Caledonia, 500–941 m, 26 October 2008. MNHN-IK-2016-2539, station DW2066 (25.283°S, 168.917°E), Athos Seamount, Norfolk Ridge, New Caledonia, 834–870 m, 26 October 2008. MNHN-IK-2011-1000, station CP31 (23.144°S, 166.859°E), south-eastern slope of New Caledonia, 850 m, 29 August 1985; all these specimens deposited in Muséum national d’Histoire naturelle in Paris, France. NIWA 82680, 36.810°S, 177.470°E, Bay of Plenty, New Zealand, 878 m, 22 April 2012; NIWA163516, ESNZ stn Z9224 (37.1°S, 177.3°E), Bay of Plenty, New Zealand, 614 m, 13 August 1998; the two specimens deposited in ESNZ National Invertebrate Collection in Wellington, New Zealand. Other material: MBM286881, isolated fragment from NIWA82680; MBM286882, isolated fragment from NIWA163516; they deposited in Marine Biological Museum, Chinese Academy of Sciences, China.

#### Diagnosis

Colony with various color after collection, including white, brown to deep ochre. Axis white to light brown. Polyps usually long tubular or club-shaped with expanded distal part, some conical or wart-like, biserially or irregularly arranged. Polyp body wall usually thick, occasionally thin and semitransparent. All sclerites with two rounded ends, occurring as slightly ridged and granulated rods in tentacles, and granulated flattened rods to thick elongate scales in polyp body wall and coenenchyme.

#### Description

Paralectotype ZMA Coel. 02711 incomplete with the holdfast not recovered (Fig. 8A). Axis white to very light brown, covered with a layer of semitransparent coenenchyme. Branches irregularly pseudo-dichotomous. Polyps tubular, about 1.2–4.7 mm long by 0.4–1.4 mm wide, with a nearly white, thin and semitransparent body wall, leading the polyps easily fall off the colony (Fig. 8E, F). Their distal part slightly expanded and became very light brown after fixation. Polyps biserially arranged on two sides of branches, occasionally arranged on the front and perpendicular to the colony plane, 1.3–4.4 mm apart. Tentacles folded over the oral disk forming a pseudo-operculum. Tegument and nematocysts absent. Other measurement data can be found in Pante et al. (2013).

Sclerites of *I. armata* densely arranged in polyps and coenenchyme, almost with two rounded ends and smooth profiles, occasionally cross-shaped. Rods in the back of the tentacle rachis usually longitudinally arranged, measuring 40–332 μm long by 10–63 μm wide (e.g., Figs. 6E, 8D). Among them, the large ones having a slightly ridged and granulated appearance, while the small ones usually non-granulated (e.g., Fig. 8H, J). Flattened rods to elongate thick scales in the polyp body wall longitudinally or obliquely arranged and with granulated surface, some of them with a slightly medial constriction, measuring 50–466 μm long by 25–119 μm wide (e.g., Fig. 8B, C, G, I, K). Sclerites in the coenenchyme same as the polyp body wall and arranged along the branch.

#### Variability in external morphology

Ten specimens (including MNHN-IK-2008-1505, 2008-1411, 2008-1504, 2009-1632, 2008-1507, 2008-1506, 2008-1510, 2016-2540, 2016-2539, 2011-1000) incomplete, cream-coloured to brown after collection (Figs. 9A, E; 11A). Their axis cream-coloured to brown and covered with thick coenenchyme. Their polyps usually large, stout and tubular with opaque and thick body wall, some of them slightly curved with expanded distal part, occasionally small and conical (Figs.9B, C, F, G, I; 11B, C), firmly attached to the branch, 1.0–5.9 mm long. Polyps usually biserially arranged, some of them present on the front while absent on the back of the colony, occasionally irregularly arranged on all sides. The surface of polyps and coenenchyme without tegument but occasionally with abundant rice-shaped nematocysts (Fig. 9J).

Specimen NIWA82680 incomplete and cream-coloured after collection (Fig. 11G). The central axis also cream-coloured and covered with thick coenenchyme. Polyps slender and tubular with expanded distal tentacular part and thick body wall (Fig. 11H), 2.0–5.5 mm long, usually biserially and alternately arranged, occasionally perpendicular to the colony plane. Polyps firmly attached to the branch. Tegument and nematocysts absent.

**Figure 11.**
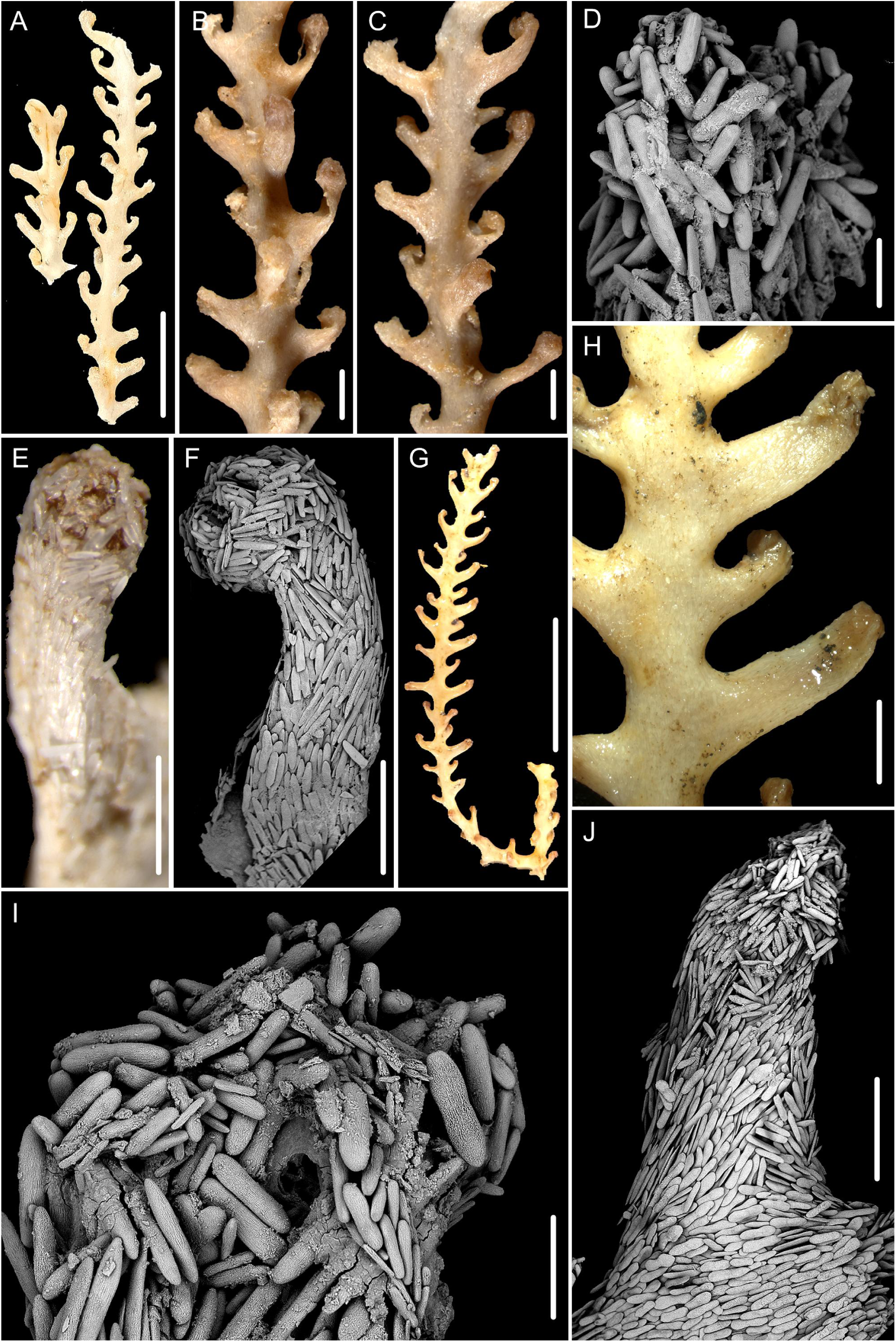
The external morphology and polyps of *I. armata* form 3, specimen MNHN-IK-2016-2539. **(A**–**F) and NIWA82680 (G**–**J).** (A, G) the separated sample; (B) the front side of branch under a light microscope; (C, H) the back side of branch under a light microscope; (E) single polyp under a light microscope; (F, J) single polyp under SEM; (D, I) tentacular part under SEM. Scale bars: 2 cm (G); 1 cm (A); 2 mm (B, C, H); 1 mm (E, F, J), 200 μm (D, I).

**Figure 12.**
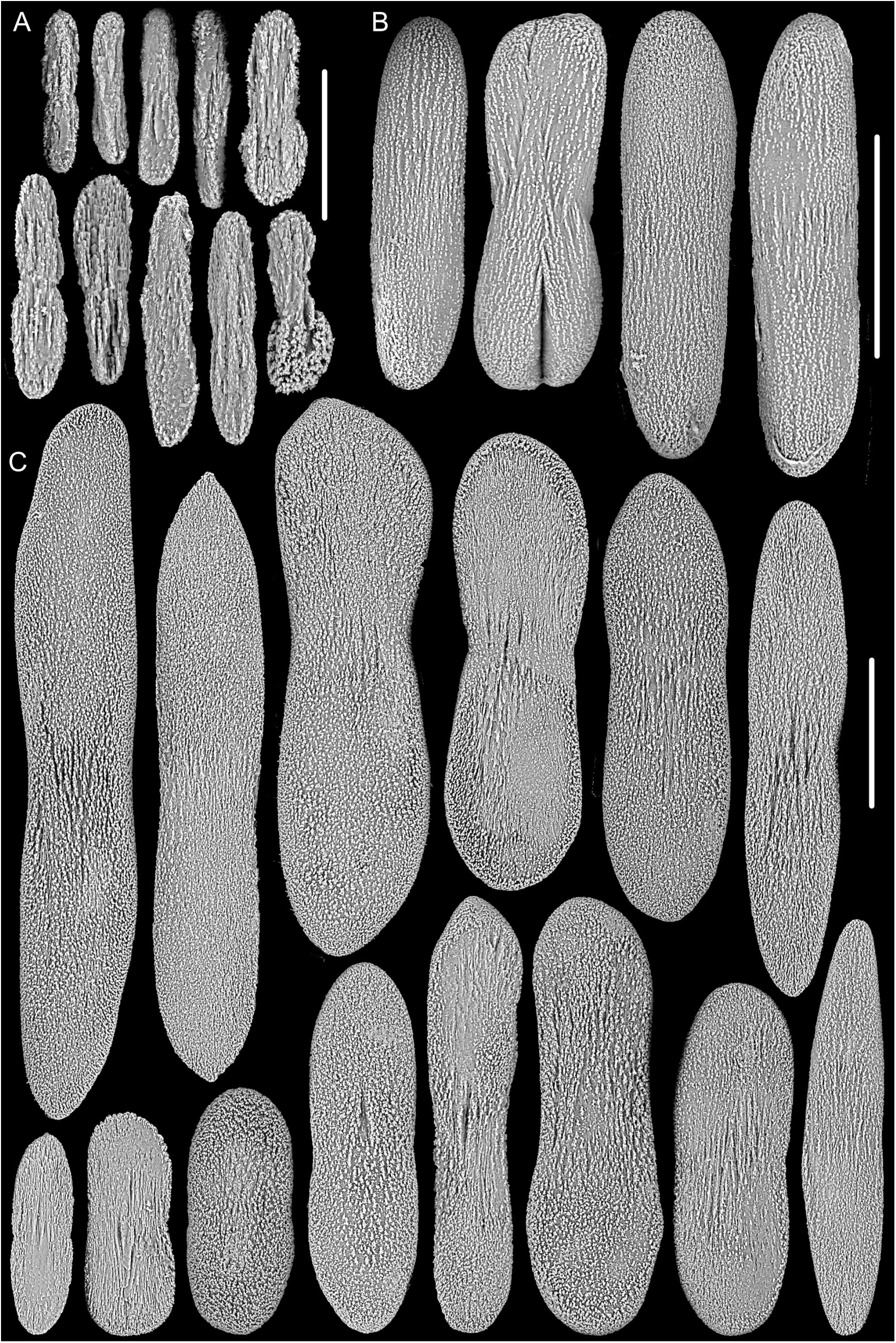
Sclerites of *I. armata* form 4, specimen NIWA82680. (A) small rods in tentacles; (B) large rods in tentacles; (C) sclerites in the polyp body wall and coenenchyme. Scale bar: 100 μm (B, C), 50 μm (A).

Specimen NIWA163516 incomplete and planar with deep ochre color after fixation, about 22 cm in height (Fig. 13A). The central axis light brown, covered with a thick layer of coenenchyme (Fig. 13B). Polyps conical and short tubular with a uniform size, 2–3 mm long, 1–2 mm wide, closely arranged on all sides of the branches (Fig. 13B, C). Polyps with an opaque and thick body wall, firmly attached the branch. Tentacles indistinct and tightly folded over the distal part of polyps, sometimes forming a pseudo-operculum. The surface of polyps and coenenchyme covered with tegument (Fig. 13B, C), a layer of thick outer coenenchyme, the structure similar to *Jasonisis thresheri* Alderslade & McFadden, 2012. The tegument packed with dense rice-shaped nematocysts, measuring 10–20 μm in length (Fig. 13E, G). The sclerites in polyp body wall and coenenchyme relatively narrow, covered with a very densely granulated appearance, some of them with irregular protrusions (Fig. 14C).

**Figure 13.**
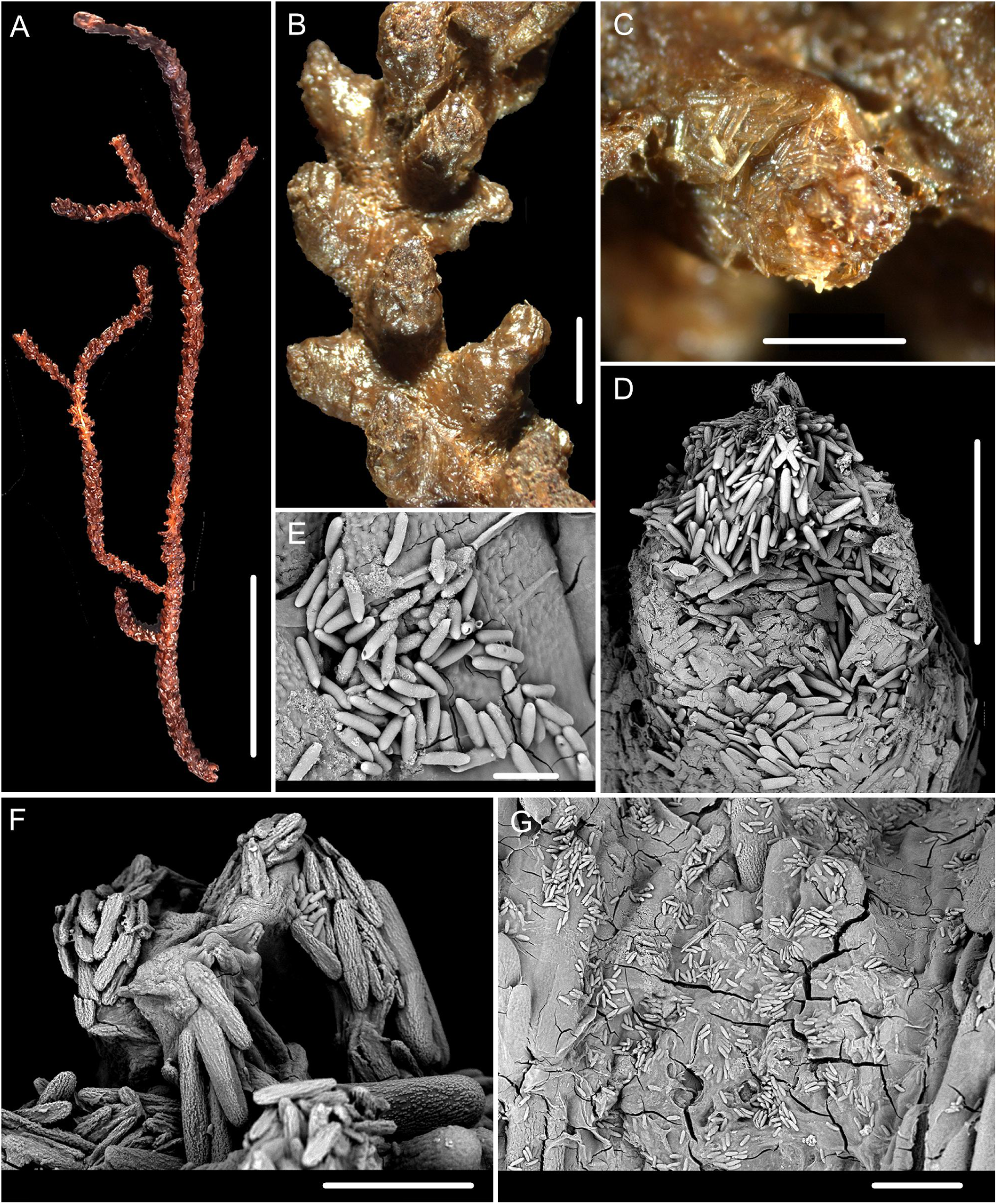
External morphology and polyps of *Isidoides armata* form 5, specimen NIWA163516. (A) specimen after fixation; (B) part of branch under a light microscope; (C, D) single polyp under a light microscope and SEM; (E, G) nematocysts on the surface of polyps and coenenchyme under SEM; (F) tentacular part under SEM. Scale bars: 5 cm (A); 2 mm (B); 1 mm (C, D); 100 μm (F, G); 20 μm (E).

**Figure. 14.**
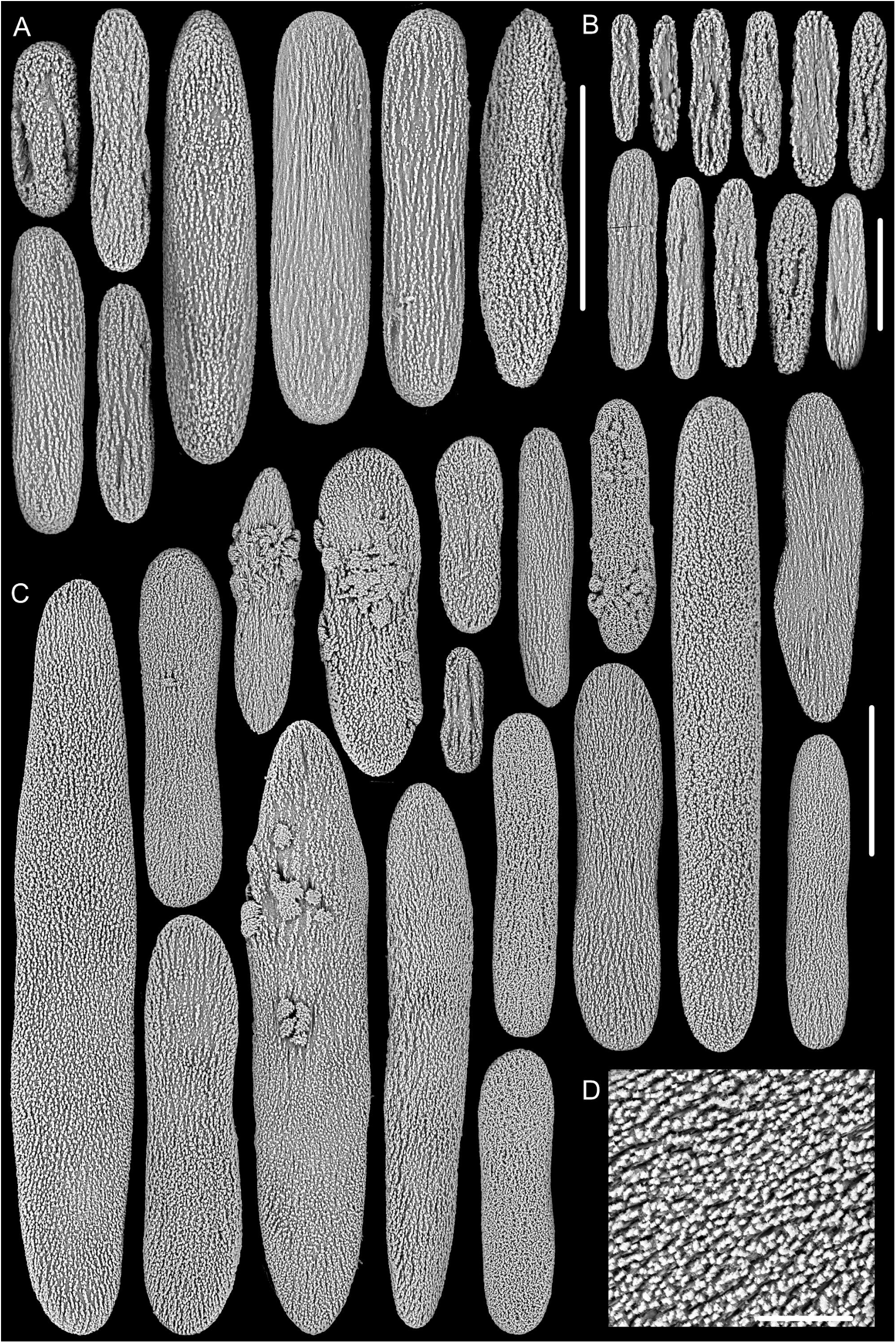
Sclerites of *Isidoides armata* form 5, specimen NIWA163516. (A) large sclerites in tentacles; (B) small sclerites in tentacles; (C) sclerites in the polyp body wall and coenenchyme, some of them with irregular protrusions on surface; (D) surface of the sclerites in polyp and coenenchyme. Scale bar: 100 μm (A, C), 100 μm (A), 50 μm (B), 20 μm (D).

#### Type locality

10.65°S, 123.67°E, south of the western tip of Timor, 520 m (Pante et al., 2013).

#### Distribution

5.9°S, 132.945°E, eastern slope of Elat, Kei Islands, Arafura Sea, 984 m (Nutting, 1910); 10.65°S, 123.67°E, south of the western tip of Timor, 520 m (Pante et al., 2013); New Caledonia to New Zealand, including south-eastern slope of New Caledonia, Norfolk Ridge, Loyalty Ridge, Kermadec Ridge and Bay of Plenty, 500–1065 m (Table 5, Pante et al., 2013, present study).

**Table 5.**
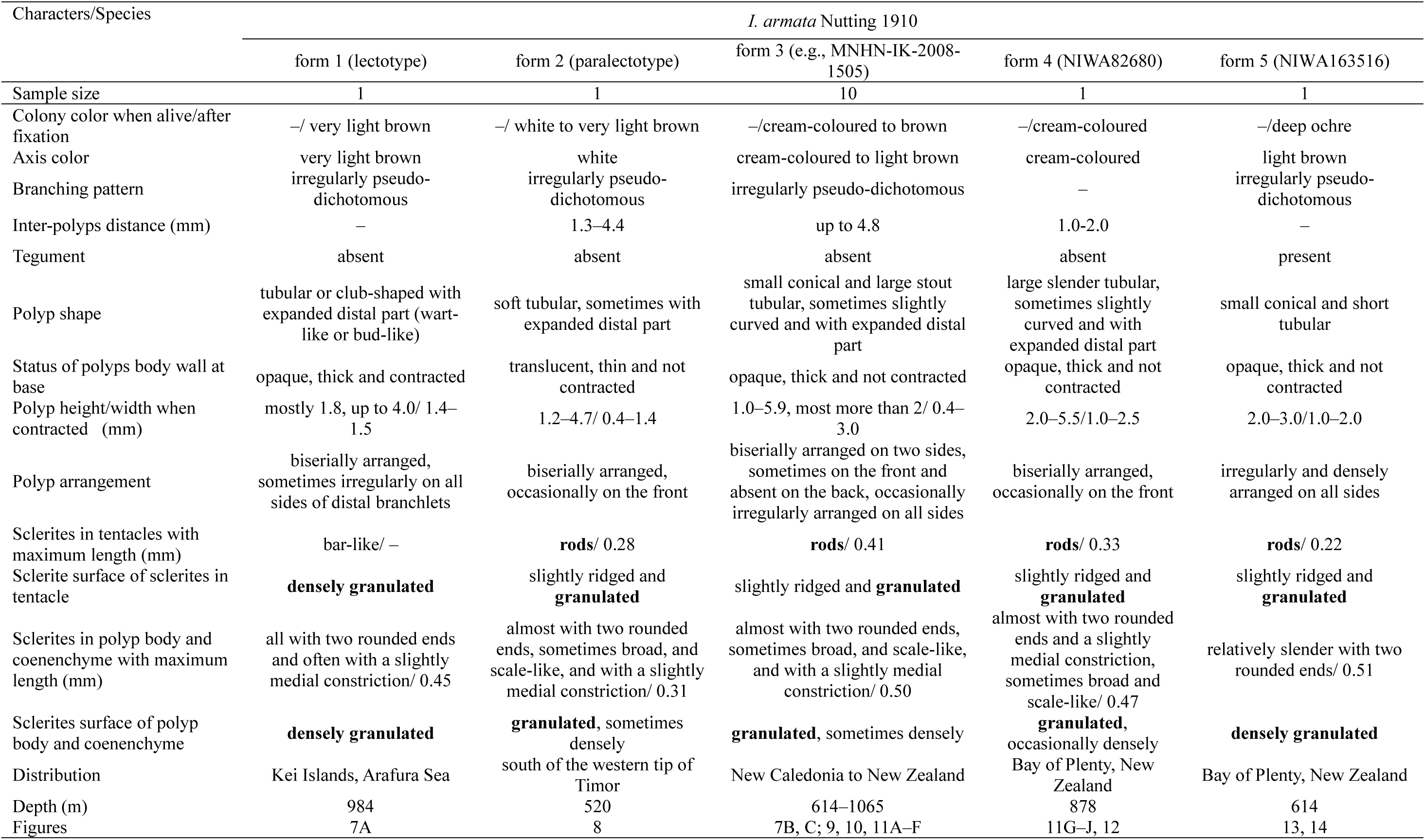
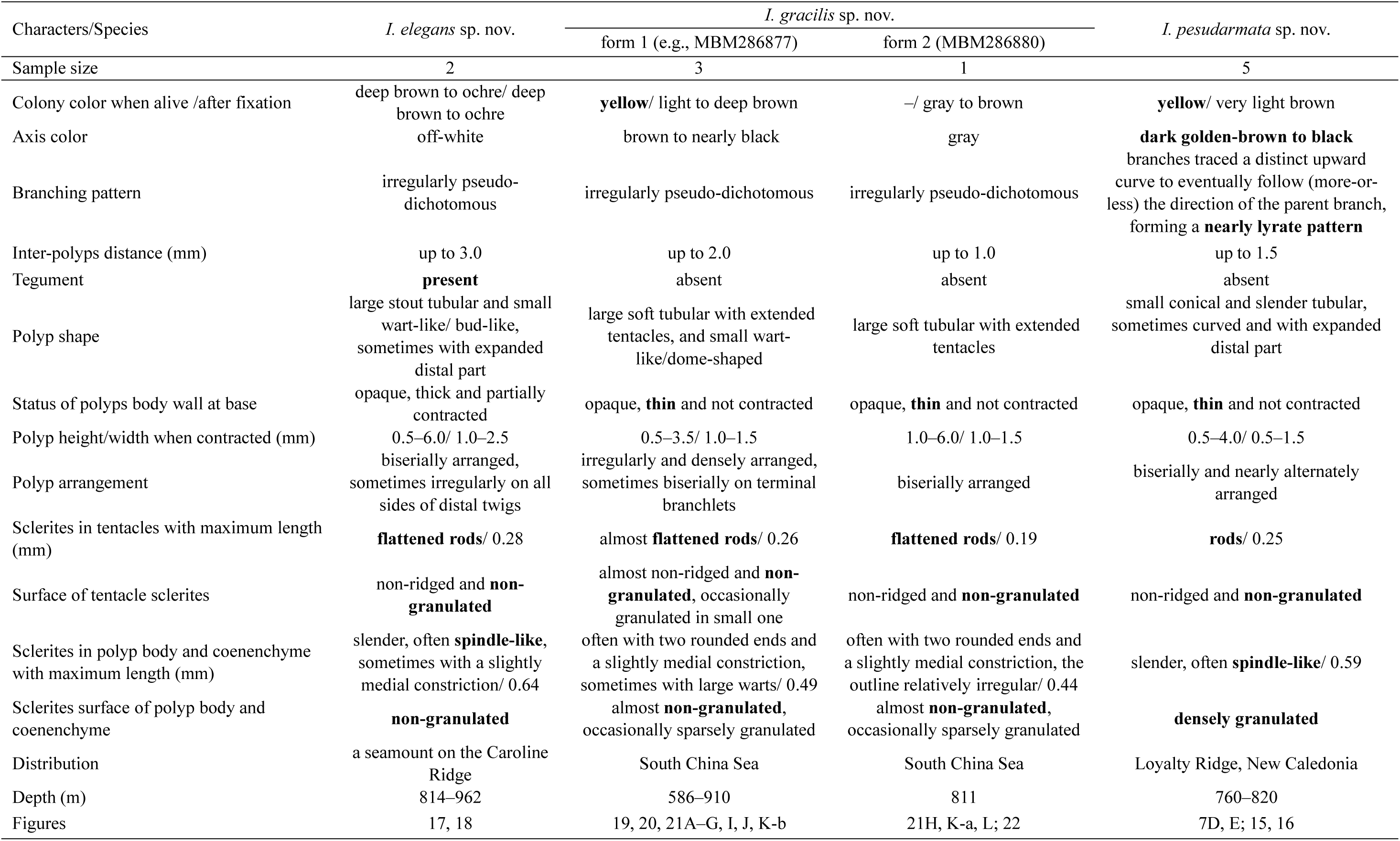
Morphological features and comparisons among *Isidoides* species. The diagnostic features for each species are presented in bold. “–” means unknown data.

#### Remarks

The known specimens of *I. armata* can be classified morphologically into five forms, namely form 1 (lectotype), form 2 (paralectotype), form 3 (specimens MNHN-IK-2008-1505, 2008-1411, 2008-1504, 2009-1632, 2008-1507, 2008-1506, 2008-1510, 2016-2540, 2016-2539, 2011-1000; AM G:16695), form 4 (specimen NIWA82680) and form 5 (specimen NIWA163516). The five forms of *I. armata* all have pseudo-dichotomous and irregular branching, light-coloured of axis, and the same sclerite forms, including slightly ridged and granulated rods in the tentacle rachis (Figs. 6D, E, G, N; 8H, J; 10B; 12B; 14B) and flattened and granulated rods to elongate thick scales (Figs. 6K, M; 8I, K; 10C; 12C; 14C) in the polyp body wall and coenenchyme. Nonetheless, obvious morphological variations were observed in colony color, polyp shape, thickness, and arrangement (Table 5). However, these differences were not constant and were possibly caused by different growth stages or environments. Given that the sclerite forms are stable and reliable for octocoral species identification (e.g., Xu et al., 2021a), we identified the five forms as the same species.

The morphological identification was also supported by the phylogenetic analysesis of sequencing data (Sanger and UCEs), which showed that distances bewteen the forms 3, 4 and 5 of *I. armata* are within intraspecific differences (see the genetic analysis aforementioned). SODA species delimitation was more nuanced, as it either lumped all forms of *I. armata* or separated the form 5, form 4 and two specimens of form 3 (MNHN-IK-2008-1507 and MNHN-IK-2016-2540) from the rest of *I. armata*. These latter two specimens, along with MNHN-IK-2008-1632, originated from the slope of New Caledonia, while all other *I. armata* specimens included in the SODA analysis were collected from adjacent seamounts. As SODA can be sensitive to population structure (Rabiee and Mirarab, 2021), we decided to include all forms of *I. armata* as one species. Finally, the haplotypes A and B defined by Pante et al. (2013) were identified as form 3, while their haplotype C is described herein as a new species, *I. pseudarmata* sp. nov.

### *Isidoides pseudarmata* Xu, Bilewitch & Pante sp. nov

Figs. 6F; 7D, E; 15; 16

urn:lsid:zoobank.org:act:729D9319-A9A8-4E51-AE8E-65B7E3660501

*Isidoides armata* haplotype C: Pante et al., 2013: 367–376, fig. 5.

#### Material examined: Holotype

MNHN-IK-2008-1869, station DW3036 (22.678°S, 168.965°E), Walpole Island, Loyalty Ridge, New Caledonia, 800 m, 16 October 2008; **Paratypes:** MNHN-IK-2008-1875, station DW3032 (22.648°S, 168.980°E), 760–820 m; MNHN-IK-2008-1867, MNHN-IK-2008-1509 and TER1062, station DW3036 (22.678°S, 168.965°E), 800 m; MNHN-IK-2008-1508, station DW3031 (22.646°S, 168.975°E), 720 m; all collected from Walpole Island, Loyalty Ridge, New Caledonia on 16 October 2008, and deposited in Muséum national d’Histoire naturelle in Paris, France.

#### Diagnosis

Colony yellow when alive and light brown after fixation. Axis dark golden-brown to black. Branches traced a distinct upward curve forming a nearly lyrate pattern. Polyps slender and tubular with expanded distal part, occasionally small and conical, biserially and alternately arranged. Polyp body wall thin but not semitransparent. Non-granulated rods in tentacles and densely granulated and slender, spindle-like flattened rods in polyps and coenenchyme.

#### Description

Specimen of holotype MNHN-IK-2008-1869 incomplete and broken, very light brown after fixation (Fig. 7E). Axis dark golden-brown to black, covered with a layer of thin coenenchyme. Branches irregularly pseudo-dichotomous branched. The terminal branchlets short, up to 11 cm long.

Polyps tubular and often curved with slightly expanded distal part, occasionally small and conical, 0.5–4.0 mm long and 1.0–1.5 mm wide (Fig. 15A, B). Polyps body wall thin but not semitransparent, and the polyps firmly attached and not easily detached from the branch. Polyps parallel to the colony plane, usually biserial and alternating between large and small forms, obliquely towards the distal end, up to 1.5 mm apart. Tegument and nematocysts absent.

**Figure 15.**
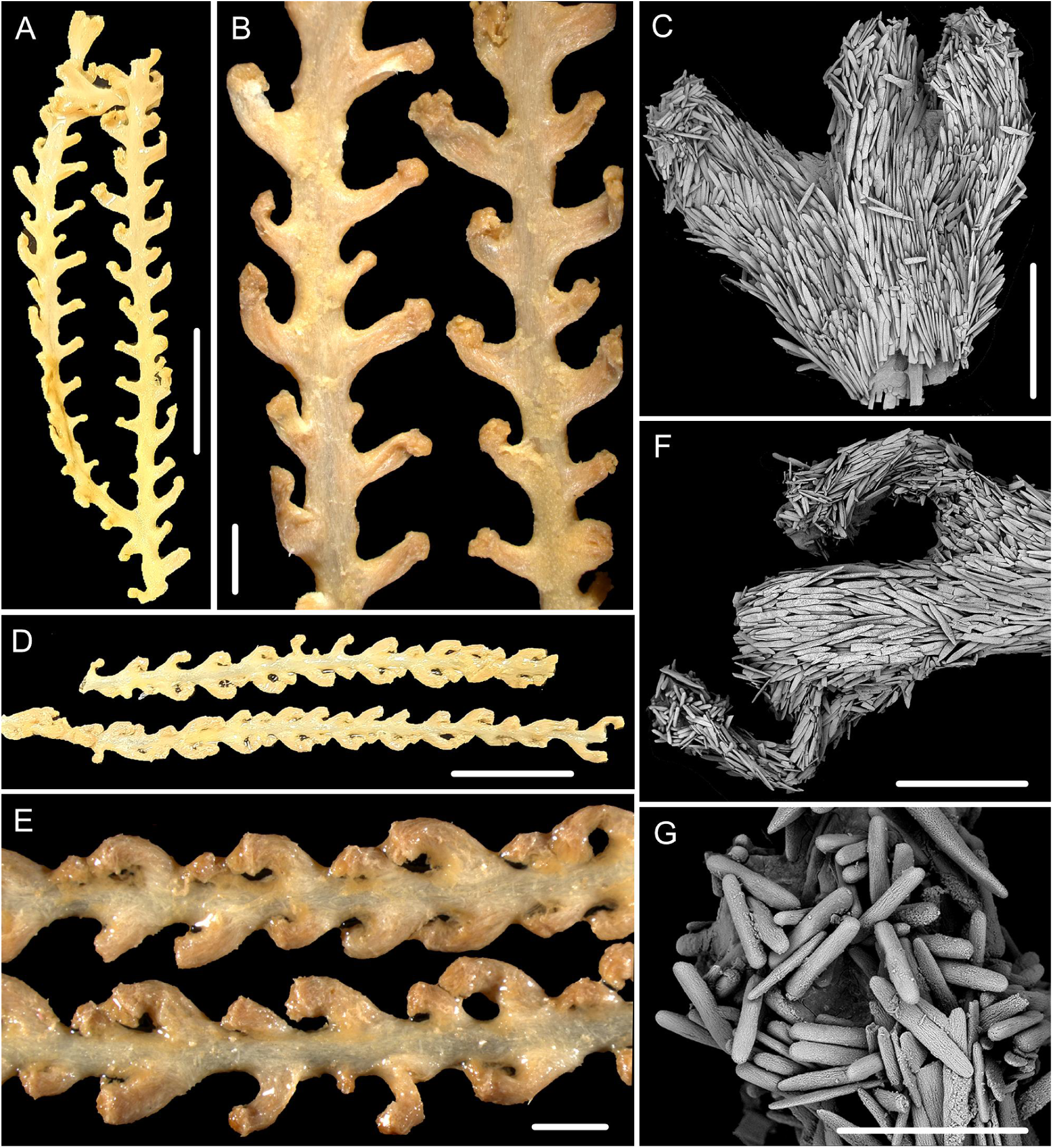
The external morphology and polyps of holotype of *I. pseudarmata*, specimen MNHN-IK-2008-1869 (A–C), and the paratype, specimen MNHN-IK-2008-1875 (D–G). (A, D) the separated sample; (B, E) two parts of branch under a light microscope; (C, F) single polyp under SEM; (G) tentacular part under SEM. Scale bars: 1 cm (A, D), 2 mm (B, E); 1 mm (C, F), 300 μm (G).

Sclerites densely arranged in polyps and coenenchyme (Figs. 15C). Rods in the back of the tentacle rachis with two rounded ends and a non-ridged and non-granulated appearance, occasionally flattened or crossed-shaped, measuring 63–250 μm long by 11–62 μm wide (Figs. 6F; 16A, B). Flattened rods in polyp body walls longitudinally or obliquely arranged, slender and often spindle-like with two narrow ends and a densely granulated appearance (Fig. 16E), occasionally crossed-shaped, measuring 123–397 μm long by 31–59 μm wide (Fig. 16C). Sclerites in coenenchyme same as the polyp body wall but with relatively larger size, measuring 143–590 μm long by 25–103 μm wide (Fig. 16D).

**Figure 16.**
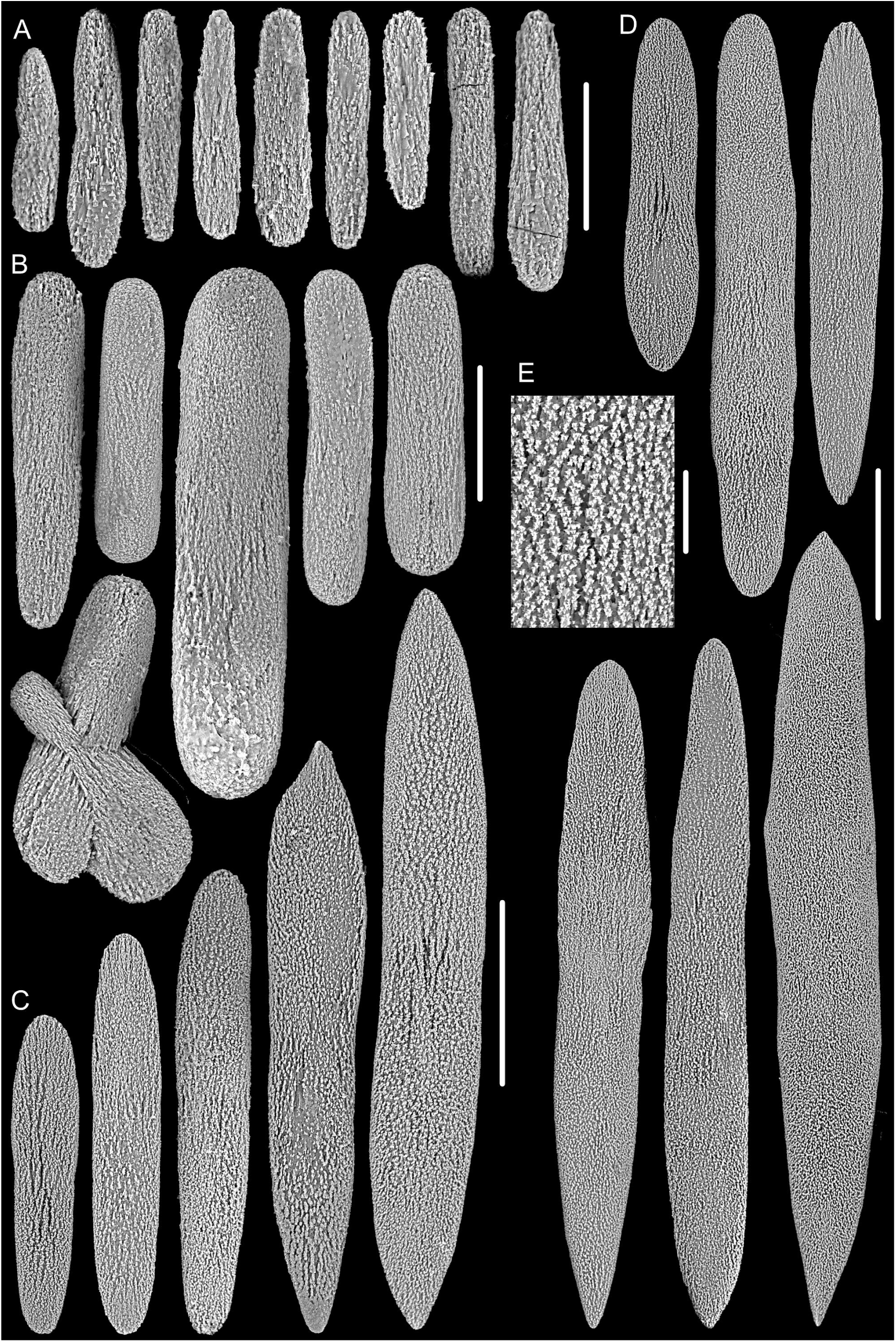
Sclerites of holotype, specimen MNHN-IK-2008-1869. (A) small sclerites in tentacles; (B) large sclerites in tentacles; (C) sclerites in the polyp body wall; (D) sclerites in coenenchyme; (E) surface of the sclerites in polyp and coenenchyme. Scale bar: 200 μm (C, D), 50 μm (A, B), 20 μm (E).

#### Variability of paratype

The colony of specimen TER1062 relatively complete and yellow after collection with the branches traced a distinct upward curve to eventually follow (more-or-less) the direction of the parent branch, forming a nearly lyrate pattern (Fig. 7D). The terminal branchlets up to 17 cm long. The polyps of other three specimens of paratype relatively small, among them, MNHN-IK-2008-1867 up to 2.5 mm, 2008-1875 and 2008-1509 up to 3.0 mm.

#### Type locality

Walpole Island, Loyalty Ridge, New Caledonia, 800 m.

#### Etymology

Composition of the Greek prefix pseud- (fake) and the species name *armata*, referring to the similarity of the new species with *I. armata*.

#### Distribution

New Caledonia with water depths of 760–820 m.

#### Remarks

The examined specimens were previously defined as haplotype C of *I. armata* (Pante et al., 2013). Based on the morphological examination and the phylogenetic analysis in this study, we separated these specimens from *I. armata* and establish a new species. *Isidoides pseudarmata* sp. nov. differs from *I. armata* by more slender sclerites in polyp body wall and coenenchyme (almost spindle-like and flattened rods, Fig. 16C, D *vs.* flattened rods to elongate thick scales, Figs. 8I, 10C, 12C, 14C), non-granulated rods in tentacles (Figs. 6O, P; 16A, B *vs.* granulated and slightly ridged, Figs. 6N, 8J, 10B, 12B, 14A), dark golden-brown to black axis (*vs.* white to light brown), relatively slender and small polyps with thin body wall (*vs.* stout and large with thick body wall) (Table 5).

### *Isidoides elegans* Xu, Bilewitch & Pante sp. nov

Figs. 6A, O; 17, 18

urn:lsid:zoobank.org:act:02A4B99F-01BC-45E8-97F3-61AF1DD01AAE

#### Material examined

**Holotype:** MBM286500, station FX-Dive 213 (10.076°N, 140.190°E), an unnamed seamount located on the Caroline Ridge, 814 m, 31 May 2019. **Paratype:** MBM286501, station FX-Dive 209 (10.078°N, 140.203°E), the same seamount on the Caroline Ridge, 962 m, 27 May 2019. The two specimens deposited in Marine Biological Museum, Chinese Academy of Sciences, China.

#### Diagnosis

Colony deep brown to ochre color when alive and after fixation with an off-white central axis. Polyps large tubular or small wart-like, usually biserially and alternately arranged, occasionally on all sides of the distal twigs. Polyp body wall thick. Polyps and coenenchyme covered by a layer of tegument packed with abundant nematocysts. Sclerites all non-granulated and flattened, including rods with two rounded ends in tentacles, and slender and spindle-like rods in polyp body wall and coenenchyme.

#### Description

Holotype planar and deep brown to ochre color when alive, about 27 cm in height and 50 cm in maximum width (Fig. 17A, C). Holdfast calcareous and nearly oval, about 35 mm in maximal length and 22 mm in width, attached firmly to a rock. The main stem below the first branch short, about 3 mm long and 4 mm in diameter at the base. The central axis solid and off-white, covered with a thick layer of coenenchyme, occasionally a calcareous but not horny bulge present on the branch axis. Branches irregularly and pseudo-dichotomously branched in one plane, and nearly parallel to the adjacent branches (Fig. 17C). Distance between adjacent branches 0.5–6.5 cm, and the terminal branchlets up to 31 cm long and 0.6–2.0 mm wide.

**Figure 17.**
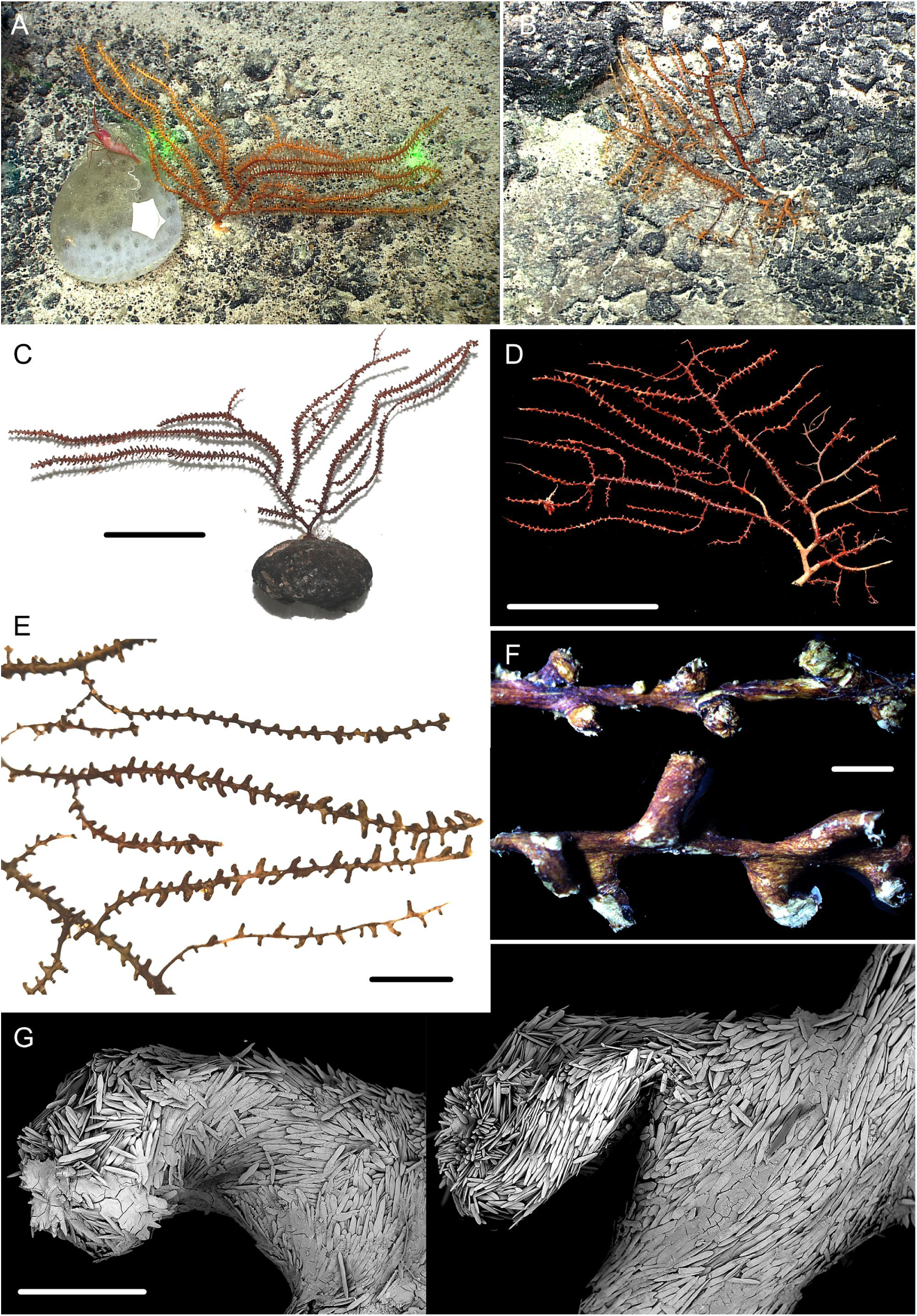
External morphology of *Isidoides elegans* sp. nov. (A, C) holotype *in situ* and after collection; (B, D) paratype *in situ* and after collection; (E) branches of the paratype after fixation; (F) two parts of branches with wart-like and tubular polyps under a light microscope; (G) tubular and wart-like polyp under SEM. Scale bars: 10 cm (C, D), 2 cm (E), 2 mm (F), 1 mm (G).

Polyps varied greatly in size and shape, including large tubular, and small wart-like/bud-like forms with expanded distal part, 0.5–6.0 mm long, 0.5–2.5 mm wide (Fig. 17F). Polyps usually biserially arranged on each side of axis, alternating between large and small forms, and occasionally present on all sides of the distal branchlets, 0.1 to 3.0 mm apart (Fig. 17C). Sometimes polyps all large tubular or all small wart-like on the branchlets (Fig. 17F). Polyps almost parallel to the colony plane, some of them curved in different directions, occasionally perpendicular to the colony plane. Polyp body wall thick and the polyps firmly attached and not easily detached from the branch. Polyps and coenenchyme covered by a thin layer of tegument (Fig. 17F), which is packed with abundant nematocysts similar to particular specimens of *I. armata*.

Polyps and coenenchyme covered with dense and irregularly arranged sclerites, all flattened and thin rods, some with a slightly medial constriction, occasionally crossed-shaped (Figs. 6A, 17G, 18). The surfaces of sclerites non-granulated, usually striped, feather-shaped or of mixed shapes, and rarely lamellar (Fig. 6H–J, L, P). Sclerites in the back of the tentacle rachis with two rounded ends, measuring 63–278 μm long by 11–50 μm wide (Fig. 18A, B). Sclerites in the polyp body wall slender and often spindle-like with two narrow to pointed ends, measuring 94–643 μm long by 25–140 μm wide (Fig. 18C). Sclerites in the coenenchyme same as the polyp body wall, measuring 136–615 μm long by 32–120 μm wide (Fig. 18C).

**Figure 18.**
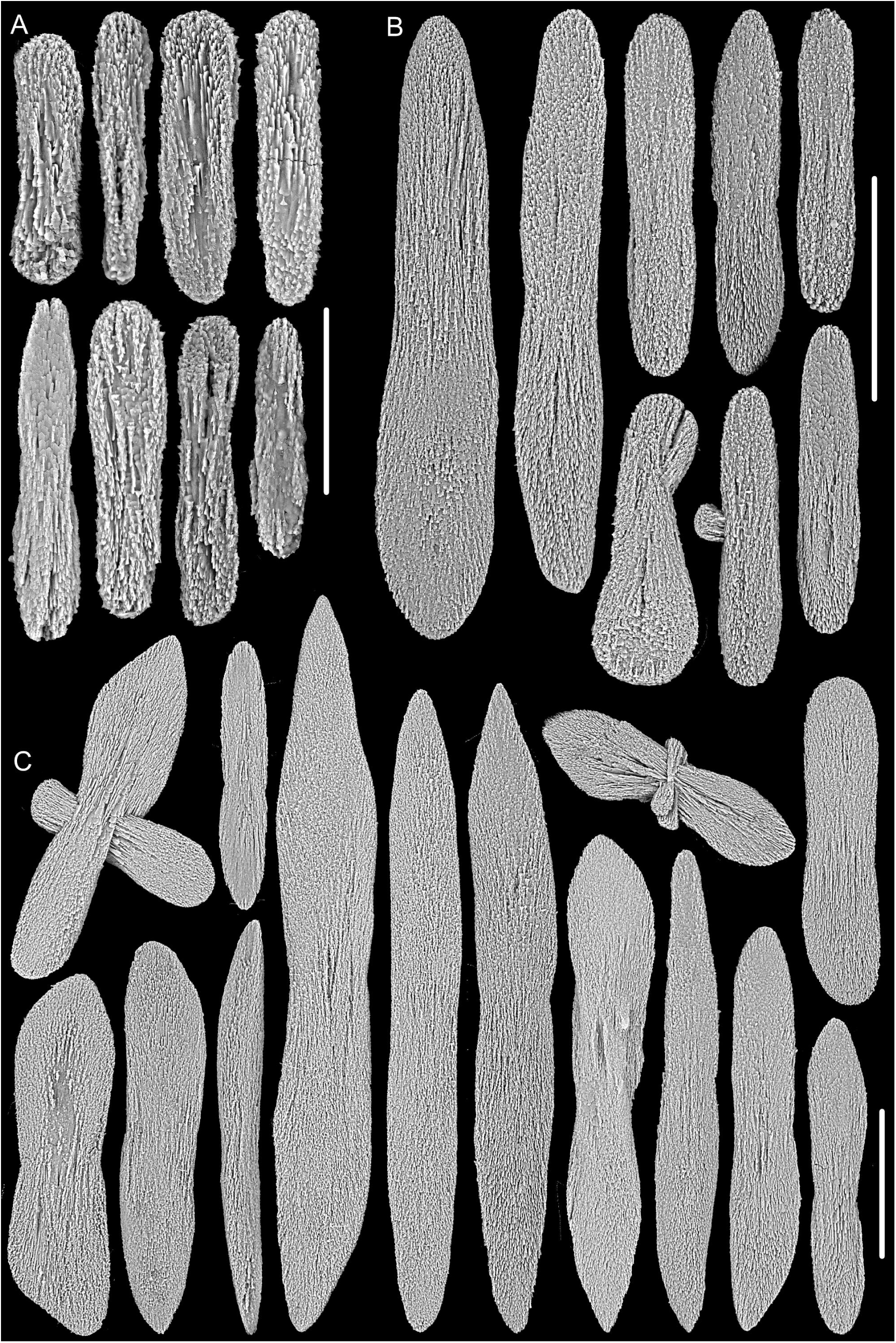
Sclerites of *Isidoides elegans* sp. nov. (A, B) small and large sclerites in tentacles; (C) sclerites in polyp body wall and coenenchyme. Scale bar: 100 μm (B, C), 50 μm (A).

#### Variability of paratype

Paratype planar with holdfast absent when it was discovered, about 15 cm high and 25 cm wide (Fig. 17B, D). The basal stem and some branches bare, with an exposed off-white axis. Main stem about 9 mm long before first branch, and 5 mm in diameter at base. Branches up to 10 cm in length and 0.6–2.0 mm in diameter. Polyps 0.5–4.0 mm long and 0.5–2.5 mm wide (Fig. 17E).

#### Type locality

An unnamed seamount located on the Caroline Ridge, 814 m.

#### Etymology

The Latin adjective *elegans* (elegant) refers to the beautiful external appearance of this species.

#### Distribution and habitat

Found on the seamounts located on the Caroline Ridge in the tropical Western Pacific with water depths of 814–962 m. The colony grows on a rocky substrate, but may still survive when detached from their holdfast (Fig. 17A, B). At the site of collection, the water temperature was 5.31–5.65°C and the salinity about 36.8 psu.

#### Remarks

*Isidoides elegans* sp. nov. is similar to *I. armata* Nutting, 1910 in having varied polyp sizes and some bud-like polyps with expanded distal part, ochre color and a layer of tegument with abundant nematocysts, but differs from the latter by non-granulated sclerites (Fig. 6H–J, L, P *vs*. granulated, Fig. 6K, M, N), flattened rods in tentacles (Figs. 6A, 18B *vs*. rods, Figs. 6D, E, G; 8H, 10B, 12B, 14A) and flattened rods in the polyp body and coenenchyme (slender and often spindle-like, Fig. 18C *vs*. elongate thick scales to flattened rods with two rounded ends, Figs. 8I, 10C, 12C, 14C) (Table 5).

### *Isidoides gracilis* Xu, Bilewitch & Pante sp. nov

Figs. 6B, C, P; 19–22

urn:lsid:zoobank.org:act:EC68E605-622F-4CB4-8FA9-13D8FCD00A6B

#### Material examined: Holotype

MBM286877, station FX-Dive 308 (15.049°N, 116.558°E), Zhenbei Seamount in the central South China Sea, 797 m, 20 July 2022. **Paratypes**: MBM286878, the Ganquan Plateau in the northwest South China Sea, 586–910 m, 14 June 2018. MBM286879, station FX-Dive 125 (22.035°N, 118.776°E), close to a submarine hydrothermal solution in the northern South China Sea, 811 m, September 2016. MBM286880, the same location and date with MBM286879, depth unknown. The four specimens deposited in Marine Biological Museum, Chinese Academy of Sciences, China.

#### Diagnosis

Colony yellow when alive and gray to brown after fixation. Axis gray to dark brown. In juveniles, polyps uniformly tubular and biserially arranged. In adults, polyps small wart-like and densely and irregularly arranged on main stem and large branches, and large tubular and biserially arranged in branchlets. Polyp body wall thin and opaque. Sclerites in tentacles, polyp body wall and coenenchyme all almost flattened and non-granulated rods with two rounded ends and a slightly medial constriction.

#### Description

The colony of holotype *in situ* planar, yellow to light brown with a large and white holdfast (Fig. 19A). The collected specimen incomplete with the holdfast not recovered, about 65 cm in height, and became light brown after fixation (Fig. 19C). Stem about 4 mm at its base, occasionally with a swollen protrusions (Fig. 19G). The central axis round and dark brown, sometimes became nearly black and a little flat. The axis of branchlets light brown, some covered with small and round pits (Fig. 21F). Branches gracile and flexible, irregularly and pseudo-dichotomously branched in one plane and without anastomosis (Fig. 19A, C). Distance between the adjacent branches 0.5–11.5 cm long, and the terminal branchlets up to 21 cm long.

**Figure 19.**
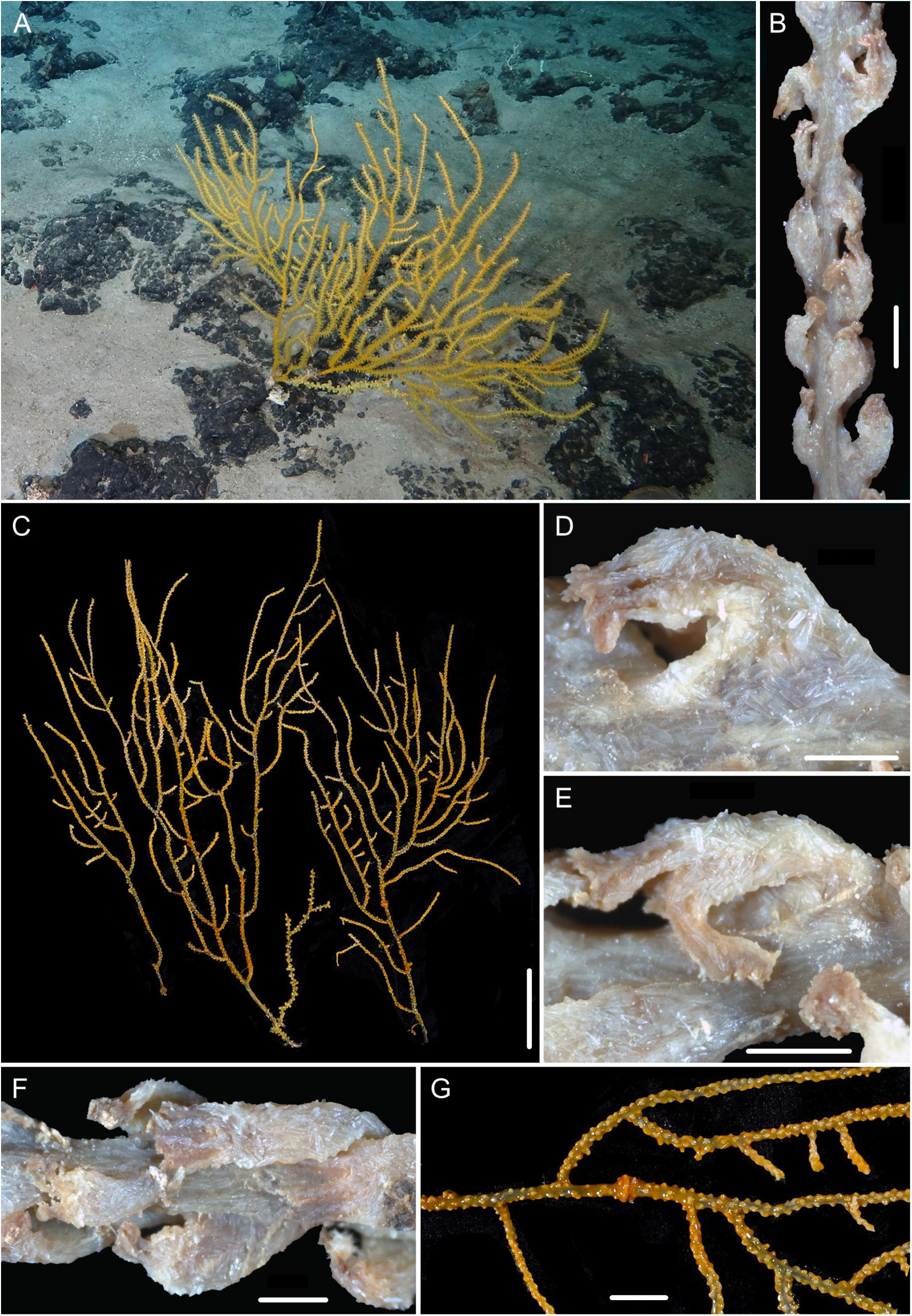
External morphology of the holotype of *Isidoides gracilis* sp. nov. form 1. (A, C) the specimen *in situ* and after collection, respectively; (B, F) part of terminal branchlet under a light microscope; (D, E) single polyps under under a light microscope; (G) part of branch with abundant wart-like polyps and a swollen protrusion. Scale bars: 10 cm (C), 2 cm (G), 2 mm (B), 1 mm (D, E, F).

Polyps vary in size and shape, including large tubular (Fig. 19B, D–F) and small wart-like (Fig. 19G), 1.0–3.5 mm long, 1.0–1.5 mm wide. Among them, small wart-like polyps irregularly and densely arranged on all sides of the main stem and large branches (Fig. 19G), while large tubular polyps irregularly arranged in the terminal branches, sometimes biserially and alternately arranged (Fig. 19B, F). Polyps with thin but opaque body wall, firmly attached and not easily detached from the branch. Tegument absent, but occasional nematocysts can be seen on the surface of polyps and coenenchyme under SEM.

Polyps, tentacles and coenenchyme covered with densely and irregularly arranged flattened rods (Fig. 21G). These sclerites are often found with a slightly medial constriction and two rounded ends, occasionally crossed-shaped or irregular or with large protrusions and grooves (Fig. 20). sclerite surface almost non-granulated, usually striped, lamellar or of mixed shapes, rarely feather-shaped, occasionally sparsely granulated in some small sclerites of tentacles (Figs. 6H, I, J, L, O; 20). Sclerites in the back of the tentacle rachis small and sometimes thick, measuring 71–262 μm long by 8–52 μm wide (Figs. 6B; 20A, B); in the polyp body wall, measuring 143–419 μm long by 22–130 μm wide (Fig. 20C). Sclerites in the coenenchyme same as the polyp body wall but occasionally relatively thick and large, measuring 113–488 μm long by 34–204 μm wide (Fig. 20C).

**Figure 20.**
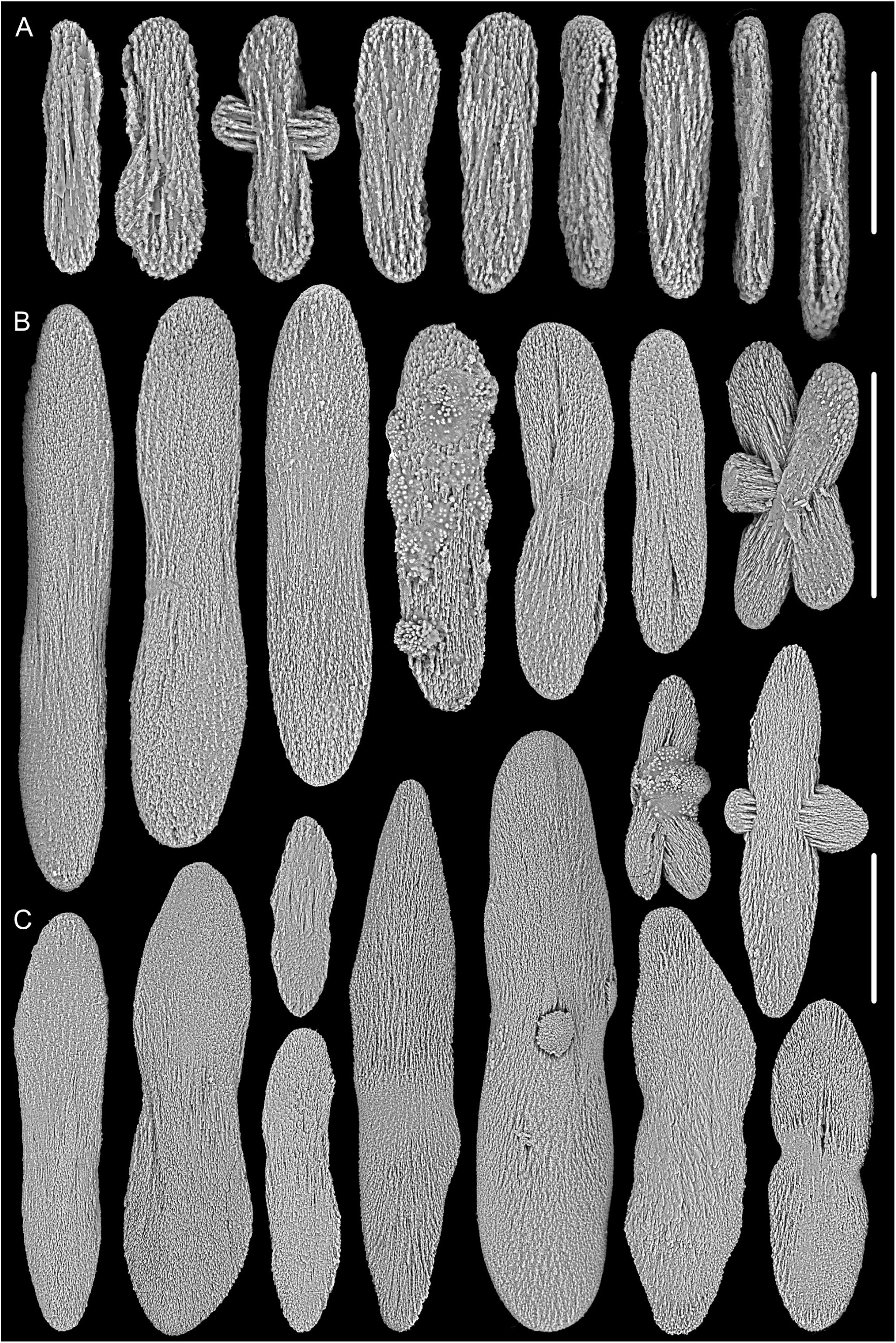
Sclerites of *Isidoides gracilis* sp. nov. form 1. (A, B) small and large sclerites in tentacles, respectively; (C) sclerites in polyp body wall and coenenchyme. Scale bar: 100 μm (B, C), 50 μm (A).

#### Variability of paratype

The colony of specimen MBM286878 incomplete and yellow after collection, about 48 cm in height (Fig. 21A). The polyps all small wart-like, irregularly and densely arranged on all sides, 0.5–1.0 mm long and up to 2 mm apart (Fig. 21B–E). The colony of specimen MBM286879 deep brown after fixation, about 65 cm in height with the terminal branchlets up to 21 cm long (Fig. 21I). The polyps 1.0–4.0 mm long and up to 1.0 mm apart (Fig. 21J, K-b). The colony of specimen MBM286880 small, about 10.5 cm in height and gray to brown after fixation (Fig. 21H). Axis gray and round. Polyps all soft and uniformly tubular with extended tentacles, 1.0–6.0 mm long, biserially arranged (Fig. 21K-a, L).

**Figure 21.**
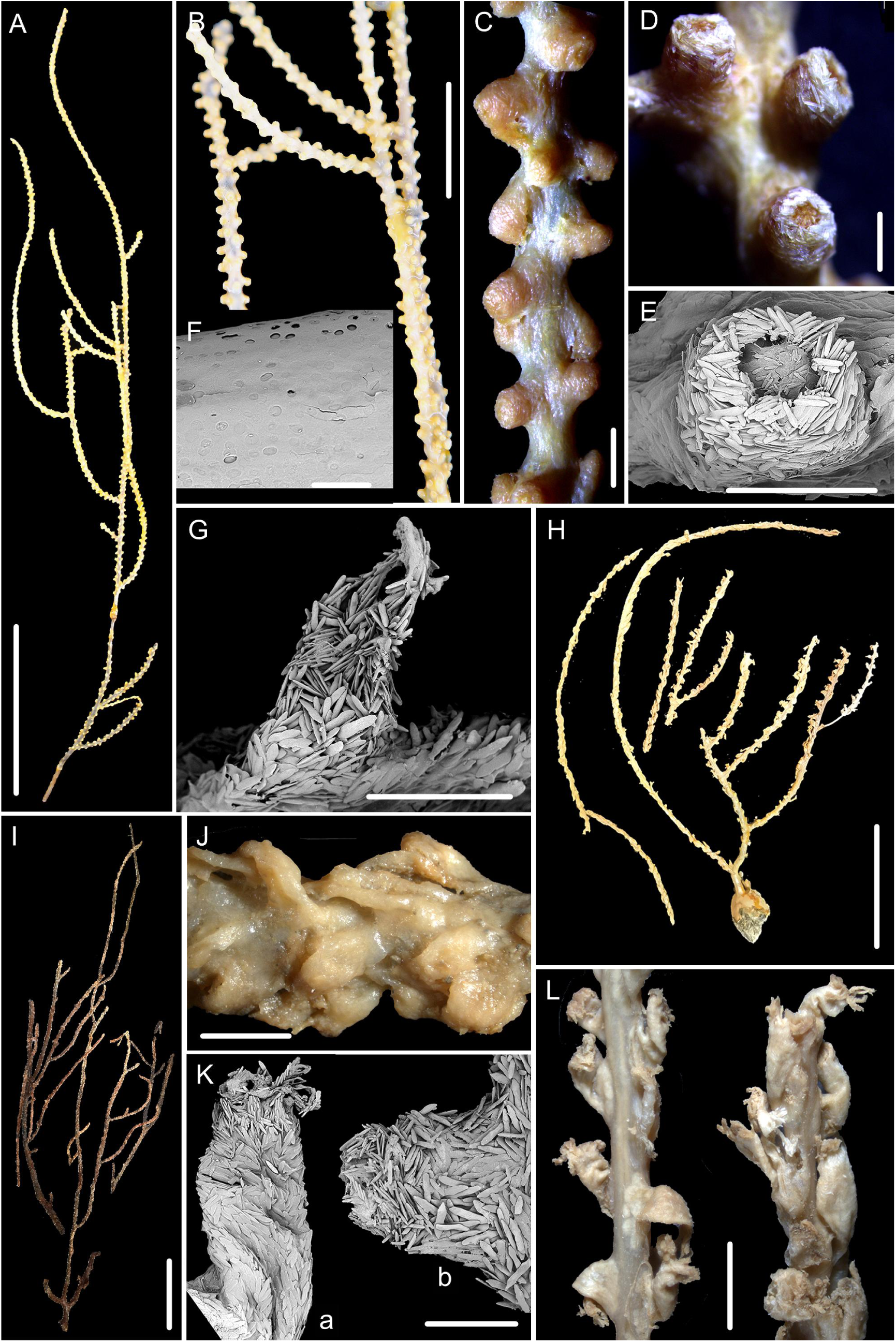
External morphology and polyps of *Isidoides gracilis* sp. nov. form 1, specimen MBM286877 (G), MBM286878 (A–E) and MBM286879 (I, J, K-b), form 2 MBM286880 (H, K-a, L). (A, H, I) the specimen after collection; (B) close-up branches; (C, D, J, K) part of branches with polyps under a light microscope; (E, G, K) single polyp under SEM; (F) surface of branch axis. Scale bars: 10 cm (A, I), 5 cm (H), 2 cm (B), 2 mm (C, J, L), 1 mm (D, E, G, K), 100 μm (F).

#### Type locality

Zhenbei Seamount in the South China Sea, 797 m.

#### Etymology

The Latin adjective *gracilis* (gracile) refers to the gracile stem and branches of this species.

#### Distribution and habitat

Found from the South China Sea at water depths of 586–910 m. Colony was attached to a rocky substrate (Fig. 19A). The water temperature was 5.85°C and the salinity about 36.3 psu.

#### Remarks

The four specimens of *I. gracilis* sp. nov. can be classified morphologically into two forms, namely form 1 (specimens MBM286878–286879) and form 2 (specimen MBM286880). They all have the same sclerite forms, characterized by a non-granulated appearance and frequently having a slightly medial constriction. While the SODA delimitation analyses suggest that forms 1 and 2 are separate species, the phylogenetic analysis of Sanger and UCE sequences indicates they fall within the range of intraspecific differences (see the genetic analysis above). Pending more data, we conservatively identified the four specimens with two forms as the same species.

Minor differences between the two forms include (1) a small colony with a round and gray axis in form 2 (specimen MBM286880), and a relatively large colony with a round and brown axis, sometimes flattened and nearly black in form 1; (2) polyps of the form 2 are uniformly tubular and biserially arranged (Fig. 21L), while the form 1 are small wart-like or large tubular, and densely arranged on each side of main stem and branches, occasionally biserially arranged on branchlets (Figs. 19B, F; 21C, J). These differences are not constant and possibly caused by different growth stages (e.g. form 2 is a juvenile and form 1 is likely an adult) or environmental conditions, and we thus treated them as conspecific variation.

*Isidoides gracilis* sp. nov. is similar to *I. pseudarmata* sp. nov. in yellow colony when alive, but differs from the latter by non-granulated and more stout sclerites in the polyp body wall and coenenchyme (Figs. 20C, 22C *vs.* granulated and slender, Fig. 16C), flattened rods in tentacles (Figs. 6B, C; 20B; 22B *vs.* rods, Figs. 6F, 16B), and presence of small wart-like and densely arranged polyps (*vs.* absence) (Table 5).

**Figure 22.**
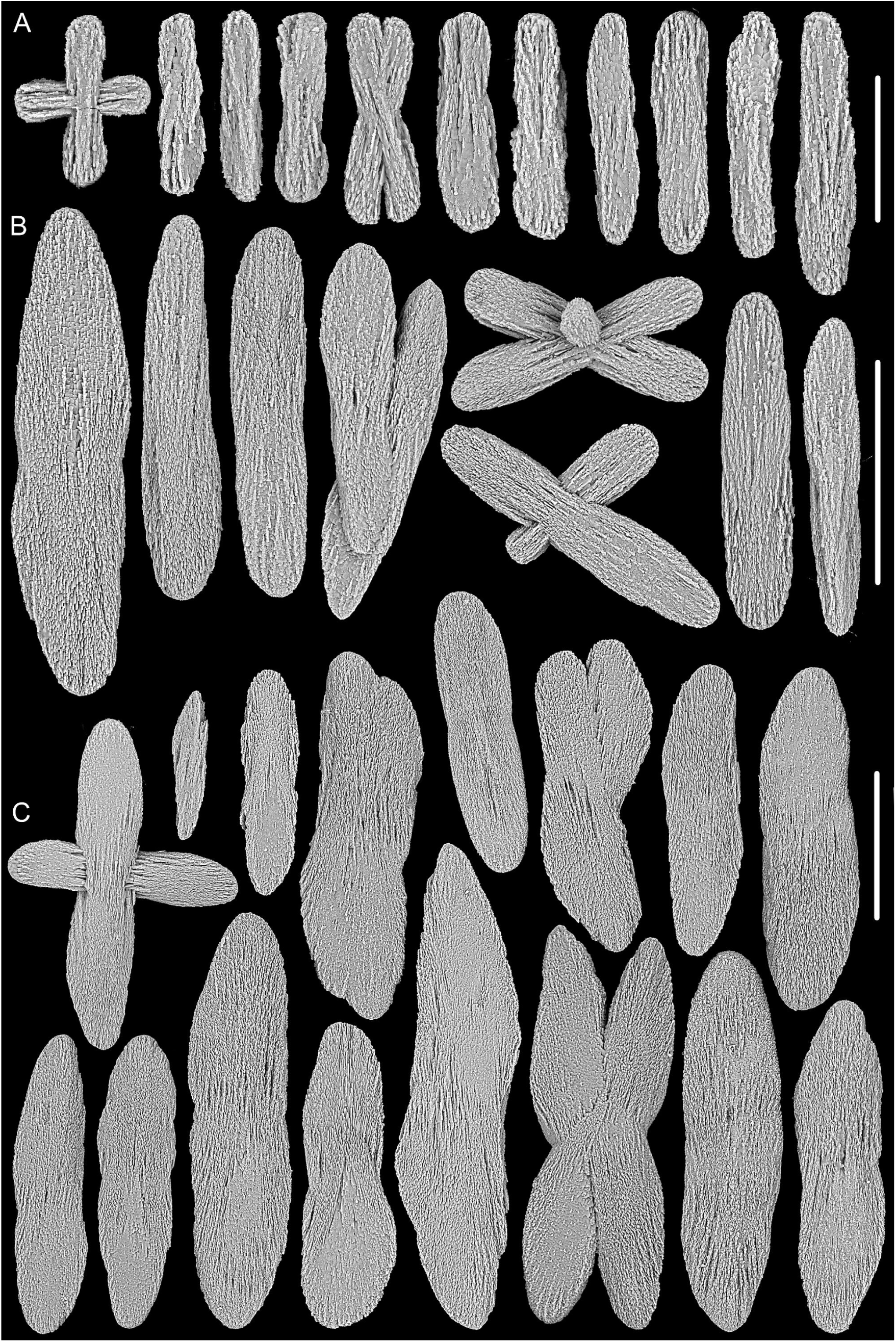
Sclerites of *Isidoides gracilis* sp. nov. form 2, specimen MBM286880. (A) small sclerites in tentacles; (B) large sclerites in tentacles; (C) sclerites in the polyp body wall and coenenchyme. Scale bar: 100 μm (B, C), 50 μm (A).

**Figure 23.**
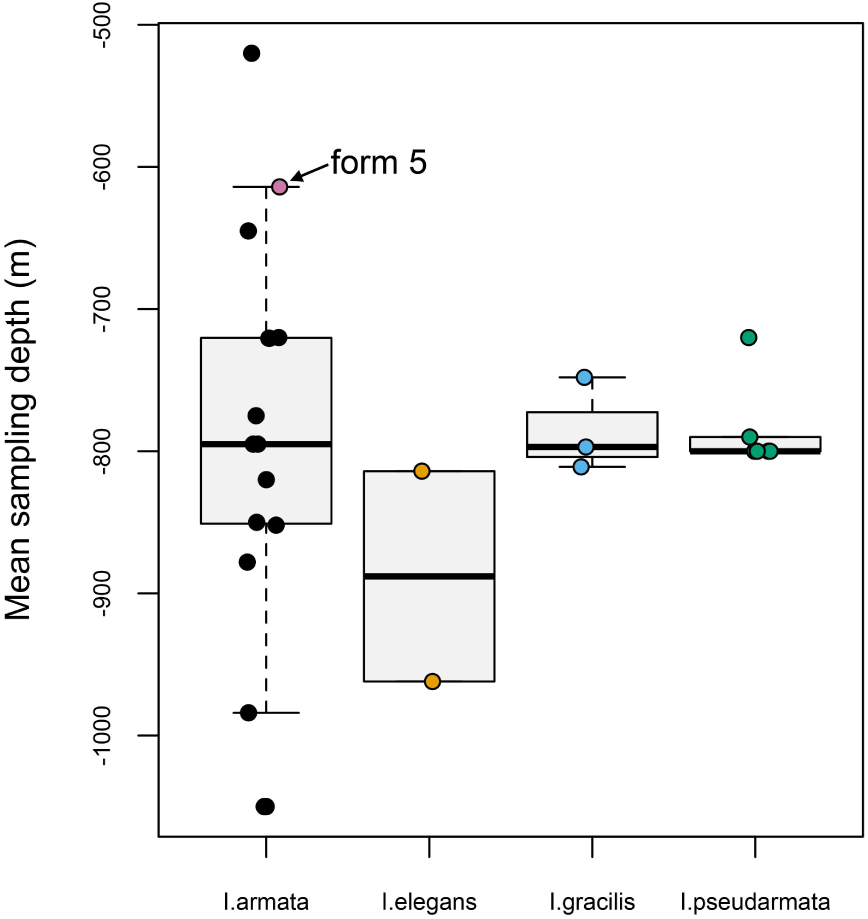
**Boxplots of sampling depths of *Isidoides* species with overlayed individual data.**

## Discussion

### High diversity of *Isidoides* Nutting, 1910

Based on available evidence, we propose four valid species of *Isidoides*, including *I. armata* Nutting, 1910, *I. elegans* sp. nov., *I. gracilis* sp. nov. and *I. pseudarmata* sp. nov. All of the known *Isidoides* specimens have been found in the Indo-West Pacific and southwest Pacific at water depths of 424–1065 m, showing an unexpected high diversity within this region (Table 1; Nutting, 1910; Pante et al., 2013; present study).

The distribution of *Isidoides* species suggests that latitude plays a greater role than depth in structuring species ranges (Figs. 1, 23). Three species—*I. elegans* sp. nov., *I. gracilis* sp. nov. and *I. pseudarmata* sp. nov. —are largely separated by latitude, whereas *I. armata*, the best-sampled species, exhibits the broadest latitudinal distribution. Its widespread range overlaps with that of *I. pseudarmata* sp. nov. in New Caledonia. In contrast, depth appears to be of secondary importance, as the bathymetric ranges of most species overlap substantially. Notably, *I. armata* stands out for its extensive distribution, both latitudinally and bathymetrically. However, these patterns must be interpreted with caution, since sample sizes remain low across all four species.

### Comparison of the phylogenetic resolution of mitochondrial genes (*mtMutS-cox1*), nuclear genes (*28S*) and ultraconserved elements (UCEs)

Recent studies have shown a higher resolving power at *28S* compared to mitochondrial markers (e.g., *mtMutS*, *cox1*) when delimiting octocoral species (e.g., McFadden et al., 2014; Quattrini et al., 2019; Xu et al., 2021a, b; Baena et al., 2024). Consistent with these findings, our study reveals that the concatenated *mtMutS-cox1* dataset only distinguishes *I. elegans* sp. nov. and fails to resolve phylogenetic relationships among all other *Isidoides* species (Fig. 2). In contrast, *28S* distinguishes four species of *Isidoides* with three of them forming a monophyletic clade respectively (Fig. 3).

The availability of data from thousands of loci results in UCEs outperforming both nuclear (*28S*) and mitochondrial (*mtMutS-cox1*) barcode genes in resolving the systematic relationships of *Isidoides* species. In this study, UCEs convincingly distinguished four monophyletic clades of *Isidoides* species with high support, including the intraspecific relationships of the three forms of *I. armata* and two forms of *I. gracilis* sp. nov. (Fig. 4 and 5). Combining the multispecies coalescent (MSP) with admixture analysis has proven useful in partitioning intraspecific clades from inter-specific ones, complementing results from phylogenetic analyses (Singhal et al., 2025). However, the combined use of phylogenomic inference and MSP remains sensitive to low taxon sampling and population structure (leading to inflated species diversity), and still benefits from complementary (morphological) evidence in cases where intra-specific variation may be mistaken for species-level differences (e.g., Erickson et al., 2021). In addition, UCEs are anchored on highly conserved genomic regions (Faircloth et al., 2012), which may lack the polymorphism necessary to reveal fine population structure.

### Important features for species delimitation of *Isidoides*

Some taxonomic characters used for the diagnosis of other octocoral taxa, such as color, polyp shape, size and arrangement, etc., are less useful for the species identification of *Isidoides* due to intraspecific morphological variations (phenotypic plasticity and/or convergent evolution). Among the specimens of *I. armata*, five morphological forms can be recognized based on colony color (white, very light brown, brown, deep ochre); polyp shape (tubular, club-shaped, wart-like, conical), thickness (opaque and thick or translucent and thin) and arrangement (biserial, or irregular); tegument (present or absent), whereas weak genetic evidence was observed for these forms (Table 5).

Variation in sclerite morphology is usually regarded as an important feature for octocoral species identification (e.g., Versluys, 1902; Bayer, 1956; Xu et al., 2021a). In the genus *Isidoides*, all species have rod-like sclerites with relatively uniform size. Only a few microstructural differences of the sclerite surface (granulated and ridged or not), the tentacle sclerites (rods or flattened rods), and the shape of polyp body and coenenchymal sclerites (often slender and spindle-like or relatively broad with two rounded ends and a slightly medial constriction) are diagnostic of interspecific differences. Based on the morphological and phylogenetic analysis of *Isidoides*, sclerites in the tentacles and surface sculpturing of sclerites (see Fig. 6) show differences and are important features for *Isidoides* species identification. However, with the exception of *I. armata*, few representative specimens of each new species were available for examination. Among the available specimens of *I. armata*, the ultrastructure of the sclerites on the surface of the polyp body and coenenchyme varied from slightly to densely granulated, according to the five recognised forms. Further sampling is required to determine how much phenotypic variation exists in these diagnostic characters for the other species, which may occur over environmental clines such as depth or latitude or in response to preservation conditions.

Based on our current diagnoses including the morphological features as mentioned above, a key is given to distinguish the *Isidoides* species:

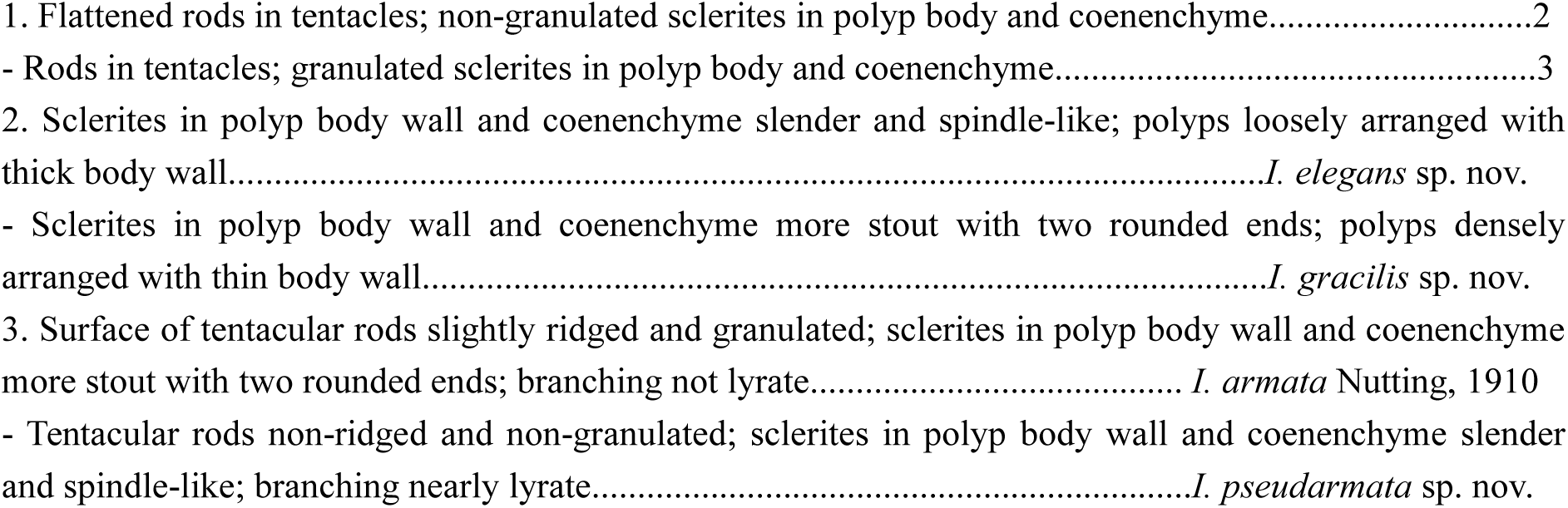

## Summary

This study described three new species and redescribed one known species of *Isidoides* Nutting, 1910, including *I. elegans* sp. nov., *I. gracilis* sp. nov., *I. pseudarmata* sp. nov. and *I. armata* Nutting, 1910. The morphological analysis showed *Isidoides* species have high intraspecific morphological variation in colony color and polyps, and the sclerite forms with their surface sculpture are important features for species identification. The phylogenetic analysis indicated that UCE phylogenomics has higher resolution than the *28S rDNA* and the mitochondrial *mtMutS* and *cox1* for species delimitation.We adopted an integrative taxonomy framework in which (1) DNA barcodes were used to test for molecular diagnosability and divergence, (2) macro- and micro-morphology was described to assess phenotypic diagnosability, (3) phylogenomic inference of gene trees and species trees were used to test for monophyly and genome-wide concordance of gene trees, and (4) species delimitation under multispecies coalescent and admixture models to test for lineage sorting and introgression among sampled and unsampled clades. These analyses hint that more diversity is yet to be revealed within the genus.

## Conclusion

Phylogenomic approaches represent a powerful means to distinguish closely related and morphologically similar species. The discovery of three new species hidden within a previously monospecific family, combined with the observed inability of single-locus barcodes to adequately distinguish these similar species indicates that, despite past molecular systematic efforts many new octocoral species await discovery in the deep-sea.

## Supporting information

Table S1. Table of detailed statistics for the UCEs pipeline

Figure S1. The structure plot (best replicate out of 20) for K=2 to 8 in the admixture analysis

## Abbreviations used in this paper include

AM: Australian Museum, Australia
BI: Bayesian Inference
ESNZ: Earth Sciences New Zealand (formerly the National Institute of Water and Atmospheric Research)
MBMCAS: Marine Biological Museum, Chinese Academy of Sciences, China ML: Maximum Likelihood
NCPOR: National Centre for Polar and Ocean Research, India
NIWA: Invertebrate Collection of the National Institute of Water and Atmospheric Research, New Zealand (now renamed the Earth Sciences New Zealand National Invertebrate Collection)
MNHN: Muséum national d’Histoire naturelle, France
SEM: Scanning Electron Microscope
TMAG: Tasmanian Museum and Art Gallery, Australia
UCEs: ultraconserved elements
USNM: United States National Museum, American
YPM: Yale Peabody Museum, American
ZMA: Zoological Museum of the University of Amsterdam, Netherlands

## Acknowledgements

We are grateful to the crew and technicians of RV KEXUE (https://cstr.cn/31114.02.MORV) for providing support and assistance in data and sample collection. We thank Dr. Jiehong Wei for his guidance on phylogenetic analysis. We also appreciate the editor for his editorial work and reviewers for their constructive comments on an early version of the manuscript. A preprint version of this article has been peer-reviewed and recommended by PCI Zool (https://doi.org/10.24072/pci.zool.100367).

New Zealand specimens collected by ESNZ during voyages KAH0011 and TAN0413 “Seamounts: their importance to fisheries and marine ecosystems” and TAN1206 “Impact of resource use on vulnerable deep-sea communities” were funded by the former New Zealand Foundation for Research, Science and Technology (CO1X0028/0224 (SFAS013/033) and CO1X0906) with additional funding from the Ministry of Fisheries (ZBD2004-01) and NOAA Satellite Operations Facility (NRAM053). An additional New Zealand specimen from station Z9224 was collected under the Scientific Observer Program funded by the New Zealand Ministry for Primary Industries (Fisheries).

The specimens hosted at the MNHN were sampled within the framework of the ‘Tropical Deep-Sea Benthos’ research program (formerly MUSORTSOM), spearheaded by the MNHN and the IRD (Bouchet et al., 2008). We are grateful to the scientists and crews of the following cruises for their sampling efforts and at-sea support: TERRASSE (R/V Alis; chief scientist Sarah Samadi; 16 Oct–31 Oct 2008; DOI: 10.17600/8100100), EXBODI (R/V Alis; chief scientists Sarah Samadi and Laure Corbari; 28 Aug–29 Sep 2011; DOI:10.17600/11100080), NORFOLK2 (R/V Alis; chief scientist Bertrand Richer de Forges; 20 Oct–06 Nov 2003; DOI: 10.17600/3100030), and BIOCAL (R/V Jean Charcot; chief scientist Claude Lévi; 08 Aug–10 Sep 1985; DOI: 10.17600/85002911). We also thank Magalie Castelin for assistance with the MNHN collections. Financial support for collection in New Caledonia during TERRASSE was provided by Sigma Xi (GIARG20061021830514629), the American Museum of Natural History (Lerner Gray Fund) and NSF (EF-0531570; EF-0531779), and the French Société Française d’Écologie. All material has been collected under appropriate collection permits and approved ethics guidelines (detailed in Pante et al., 2012).

## Supplementary

Table S1. Table of detailed statistics for the UCEs pipeline.

Figure S1. The structure plot (best replicate out of 20) for K=2 to 8 in the admixture analysis

