## Supplementary figures and images for "Unexpected diversity of *Isidoides* (Anthozoa: Octocorallia: Isidoidae) revealed by morphology and phylogenomic analysis with descriptions of three new species"

### Figure S1. The structure plot (best replicate out of 20) for K=2 to 8 in the admixture analysis

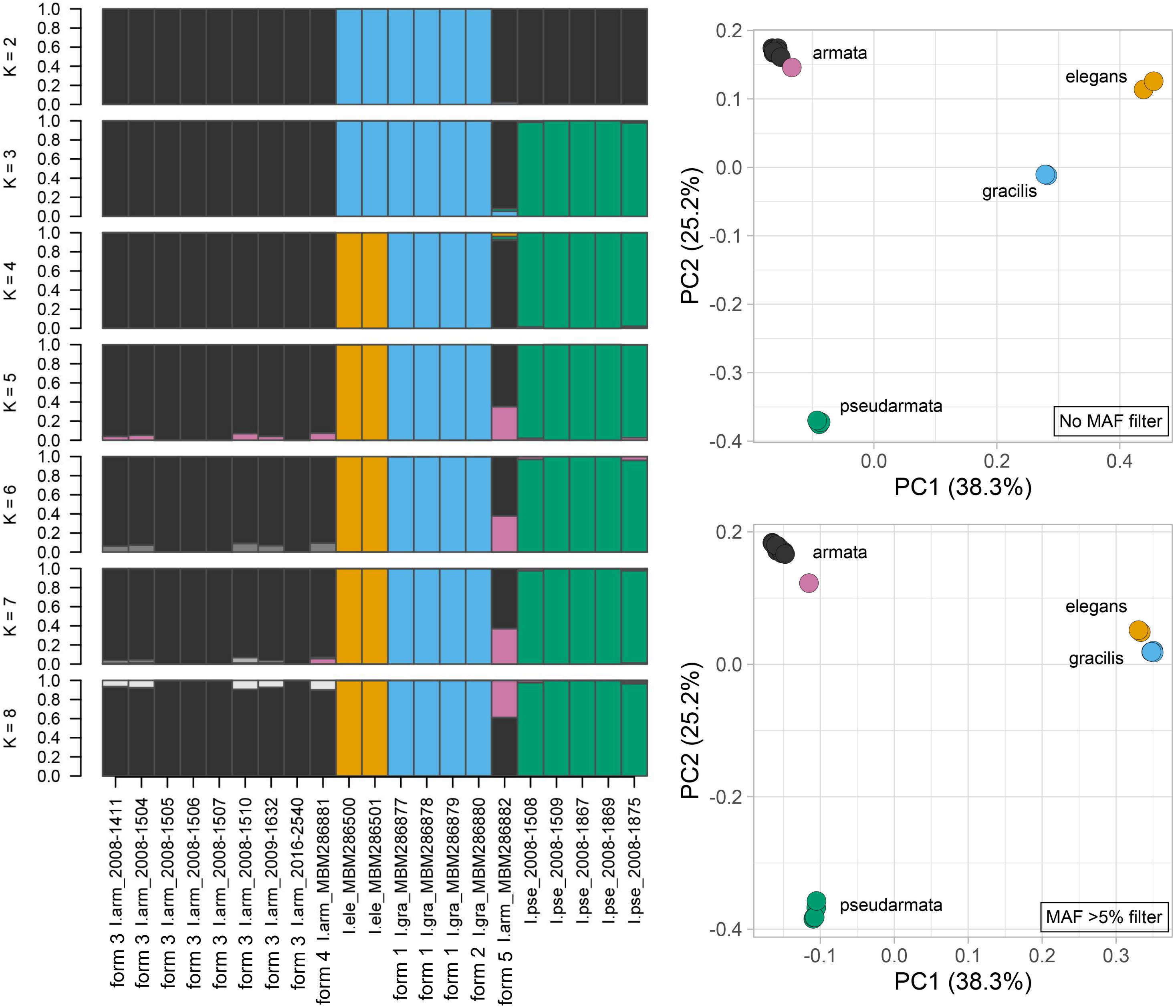
